# Crystal interface mechanics of cold-water coral skeletons

**DOI:** 10.1101/2025.10.29.685320

**Authors:** Nikolai Kvashin, Ali Ozel, Uwe Wolfram

## Abstract

Cold-water coral (CWC) skeletons utilise aragonite crystals, which possess exceptional strength (4–6 GPa), to construct a composite skeletal framework that is significantly less strong (~0.5 GPa) yet remarkably resilient. The interfacial processes governing this transition from strong crystal to tough composite remain unclear, suggesting that the organic interfaces between crystals, rather than crystal defects, control the macroscopic mechanical behaviour. We systematise the investigation of this phenomenon by employing Molecular Dynamics simulations, which reproduce experimental elastic constants within 5%. We characterise three limiting interface compositions: dry twin aragonite boundaries, hydrated protein-mediated interfaces, and water interfaces - by applying tensile and shear loading and quantifying their mechanical competence using a three-dimensional failure criterion.

Interface composition determines the mechanical hierarchy, spanning nearly an order of magnitude in strength. Dry twin boundaries provide the highest strength, reaching up to 6.5 GPa, through direct crystalline bonding. Conversely, protein-mediated interfaces exhibit the lowest strength (0.5–0.7 GPa) but demonstrate high damage tolerance. Water-mediated hydrogen-bonding networks enable this progressive failure, dissipating energy through sequential bond rupture and reformation rather than catastrophic separation. Water interfaces show thickness-dependent compliance: thin layers (<5 *Å*) retain partial electrostatic coupling (>3 GPa), while thick layers allow controlled sliding (~0.8 GPa).

These quantitative structure-property relationships provide transferable parameters for multiscale coral modelling, enabling researchers to bridge atomistic mechanisms with mesoscale mechanical response. The findings reveal how skeletal hierarchies integrate strength, stiffness, and energy dissipation, offering potential design principles for biomimetic composites that reproduce the tunable mechanical properties of biomineralised materials.

**Statement of Significance:** Cold-water coral skeletons are built from strong aragonite crystals but fail at much lower stresses than the crystals themselves. We used computer simulations at atomistic scale to understand why. Our study reveals that the interfaces between crystals, rather than the crystals control skeleton strength. We identified three interface types with different behaviours: crystalline twins are strongest and accommodate deformation through boundary migration, proteins with water layers are weaker but absorb energy through progressive failure, and water layers provide tunable properties depending on thickness. These findings explain how natural materials combine strong building blocks with compliant interfaces to achieve toughness. Our quantitative measurements provide design parameters for both predicting coral resilience to environmental stress and creating synthetic materials inspired by this hierarchical architecture.

## 1. Introduction

Cold-water corals (CWC) are abundant reef builders inhabiting cold waters at 50-4,000 m depth worldwide [1–3]. These organisms use aragonite, their primary calcium carbonate polymorph, to create extensive skeletal frameworks that host high biodiversity and provide critical ecosystem services, including carbon sequestration [4]. However, ocean acidification and climate change directly threaten CWC reefs [1–3, 5]. Climate change reduces growth of live corals [6] while accelerating dissolution of dead skeletons [7], thereby lowering net carbonate accumulation in both tropical and cold-water reef systems [6, 8, 9]. CWC reefs face particularly acute vulnerability because habitat complexity and biodiversity depend primarily on dead coral framework rather than living tissue. This dependence becomes critical as the aragonite saturation horizon, i.e., the depth below which aragonite dissolves, rises with ocean acidification. This horizon is projected to reach 70% of CWC reefs this century [2, 10]. Currently, most CWC reefs remain above this horizon (Ω_*Arag*_ >1) and maintain structural integrity [5], whereas reefs already below it (Ω_*Arag*_ <1) show reduced dead coral retention and low habitat complexity [2].

The resilience of CWC skeletons to both acidification and mechanical stressors derives from the hierarchical structure and material properties of aragonite [11–13] (Figure 1(a–f)). At the single-crystal level, aragonite exhibits exceptional stiffness (*E* = 82–138 GPa; Table 1) and strength (4–6 GPa) [11, 14–19]. However, biomineralized polycrystalline aragonite in coral skeletons shows substantially reduced properties: experimental measurements report stiffness of 45–70 GPa and strength of approximately 460 MPa [11]. Notably, future ocean conditions alter skeletal stiffness but preserve strength [11], suggesting that strength-controlling mechanisms differ from those governing elastic response. Understanding this mechanical robustness under environmental stress has direct implications for both coral conservation strategies [1, 2, 11] and the design of biomimetic materials that replicate coral skeletal architectures.

**Table 1:** Stiffness tensor for pure aragonite.

| Source | $C_{11}$ | $C_{22}$ | $C_{33}$ | $C_{44}$ | $C_{55}$ | $C_{66}$ | $C_{12}$ | $C_{23}$ | $C_{31}$ |
| --- | --- | --- | --- | --- | --- | --- | --- | --- | --- |
| Experimental values in GPa |  |  |  |  |  |  |  |  |  |
| Voigt [35] | 159.6 | 87.0 | 85.0 | 41.3 | 25.6 | 42.7 | 36.6 | 15.9 | 2.0 |
| Median from [11] | 171.1 | 110.1 | 85.0 | 41.3 | 25.6 | 42.7 | 60.3 | 18.9 | 10.6 |
| Computational values in GPa |  |  |  |  |  |  |  |  |  |
| Original Pavese [36] | 157.7 | 100.9 | 68.3 | 36.0 | 24.1 | 41.1 | 58.0 | 50.4 | 34.1 |
| Pavese based [11] | 156.1 | 96.0 | 66.5 | 34.1 | 24.6 | 38.7 | 59.1 | 50.8 | 36.1 |
| This work Pavese based | 171.8 | 106.7 | 84.2 | 42.1 | 31.1 | 46.6 | 57.5 | 46.9 | 30.2 |

**Figure 1:**
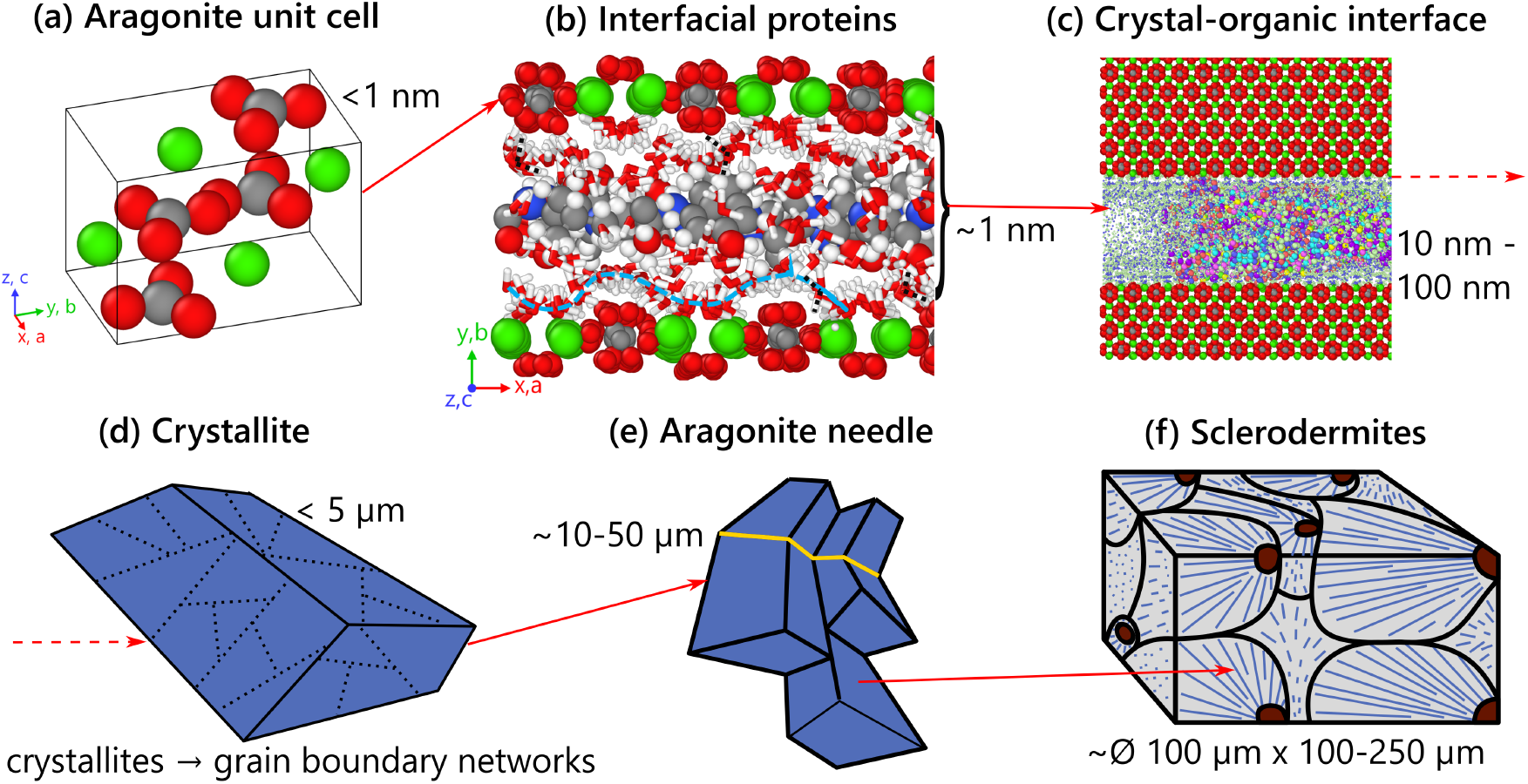
Hierarchical organization of cold-water coral skeletal structures across spatial scales (a) Aragonite unit cell (<1 nm). Atoms are shown as coloured spheres: Ca - green, C - grey, and O - red. (b) Interfacial proteins at grain boundaries (~1-100 nm). (c) Crystal-organic interface (10-100 nm). (d) Polycrystalline structure with grain boundary networks (5 µm). (e) Aragonite needle (10-50 µm). (f) Sclerodermites (∅ 100 µm *×* 100-250 µm)

At the mesoscale, CWC skeletons consist of aragonite crystals separated by organic interfaces. These anisotropic crystals organize into mesostructures that generate complex architectures spanning multiple length scales (Figure 1). Their material complexity reflects several interacting factors: crystallographic textures with preferred orientations and twin boundaries [20]; interfaces containing variable protein compositions and water content [21–26]; distributions of porosity and grain size [11, 27, 28]; and anisotropic, failure-mode-dependent mechanical responses that vary with both structural scale and loading conditions [29, 30]. Together, these features govern the transition from single-crystal properties to the effective mechanical behaviour of the polycrystalline skeleton.

At the molecular level (1-10 nm), individual aragonite unit cells with dimensions *a* = 4.961*Å, b* = 7.967*Å, c* = 5.740*Å* [31] serve as anisotropic building blocks (Figure 1(a)). The nanoscale regime (10-100 nm) features coral acid-rich proteins (CARPs) that mediate crystal nucleation and growth [20]; interfaces containing variable protein compositions and water content (Figure 1(b)). These proteins produce nanometre-thick organic interfaces between adjacent aragonite crystals within the polycrystalline matrix [21, 22, 24] (Figure 1(c)). Within these interfacial regions, water-mediated hydrogen-bond networks provide both adhesion and mechanical compliance [32, 33].

From 100 nm to tens of micrometers, crystallites coalesce into grains with complex boundary networks (Figure 1(d)). Electron backscatter diffraction (EBSD) reveals ubiquitous twins with characteristic misorientation angles of 56.3^°^and 63.8^°^, corresponding to < 001 > {310} and < 001 > {110} twin systems, respectively [20]. At the scale of 1-50 µm, aragonite crystallites approximately 5 µm in size organize into elongated needles 10-50 µm in length with preferred crystallographic orientations [20, 23, 34] (Figure 1(e)). These needles subsequently assemble into sclerodermite units that form the polycrystalline skeletal matrix [34] (Figure 1(f)). This hierarchical organization featuring twins, grain boundary networks, organic interfaces, and porosity governs the transition from single-crystal to polycrystalline mechanical behaviour. Specifically, this hierarchy explains how bulk skeletal properties (45-70 GPa stiffness, 460 MPa strength) fall substantially below single-crystal values (82-138 GPa stiffness, 4-6 GPa strength) [11, 35].

The adhesion and deformation mechanisms governing this scale transition from single-crystal properties (Table 1) to bulk skeletal properties remain unclear. Porosity alone cannot account for this mechanical property reduction [11]. Moreover, previous work shows that while ocean acidification reduces polycrystalline stiffness, it preserves polycrystalline strength [11], suggesting that different microstructural features control these two properties. This observation indicates that strength depends critically on interface characteristics rather than bulk crystal integrity or porosity.

Given this interface-dominated behaviour, we must identify which interface types control mechan-ical properties. Transmission electron microscopy (TEM) reveals grain boundary and interfacial networks that are no more than a few nanometres thick [20, 37, 38]. Based on this observed thickness, we infer that these interfaces contain primarily organic material rather than amorphous CaCO_3_ phases, which would require larger thickness. This constraint reduces possible interface compositions to three limiting cases: (i) twin boundaries providing direct aragonite-aragonite contact with specific crystallographic misorientations; (ii) organic interfaces comprising protein-mediated boundaries with CARPs and structured water; and (iii) water interfaces consisting of pure water layers between aragonite surfaces. These three interface types represent the structural extremes likely present in coral skeletons.

Previous computational work has quantified porosity effects at micro- and mesoscales [11] and identified critical links between skeletal morphology and mechanical performance [39]. However, direct experimental characterization of coral interfaces remains challenging. While nanoindentation can resolve local elastic properties [14, 17, 40], it cannot measure interface-specific behaviour across the 1 nm-1 µm regime where protein-mineral interactions dominate, nor can it quantify strength across these interfaces. Consequently, studies of aragonite-organic interfaces have focused primarily on interaction forces and elastic properties [41, 42], with few measurements of interface strength available [43, 44].

Experiments reveal complex interface morphologies and compositions [20, 37, 45], but systematic testing with controlled geometries remains challenging at these small length scales (Figure 1). Molecular Dynamics (MD) simulations overcome this limitation by enabling systematic investigation of interfacial mechanics and their contributions to stiffness, strength, and nanoscale interactions. While computational studies have examined crystalline aragonite [36, 46, 47] and larger-scale phenomena such as ocean acidification effects on skeletal density [2], most prior work emphasizes elastic properties rather than strength and failure mechanisms [17, 48, 49]. Density Functional Theory studies report comprehensive elastic constants [48], and atomistic simulations characterize surface energies and morphology [49], but neither approach directly addresses interface fracture. Understanding the strength reduction from single-crystal (4-6 GPa) to polycrystalline coral skeletons (~460 MPa) requires investigating interface failure beyond the small-strain elastic regime. Since interface strength dictates skeletal failure, investigating these mechanisms is essential for predicting CWC resilience under environmental stress.

This study investigates interface mechanics in three limiting cases, namely, dry aragonite twins, hydrated protein-mineral interfaces (aragonite + proteins + water), and pure water interfaces, to identify the mechanisms governing strength reduction in coral skeletons. We pursue four specific objectives: (i) quantify the mechanical response of these representative interface geometries under tension and shear; (ii) determine the energetic driving forces for protein-mineral adhesion in aqueous environments; (iii) explore how binding mechanisms evolve during mechanical loading and identify the factors controlling interface failure; and (iv) establish relationships between interface composition, binding strength, and mechanical performance. By combining atomistic simulations of single-crystal aragonite with systematic characterization of these limiting interface types, we identify how interfaces govern the transition from crystal-scale properties to bulk skeletal behaviour.

## 2. Methods

To investigate relevant interface systems, we build on previous studies which have characterized the stiffness and strength of single crystal aragonite [11, 16, 17, 50, 51]. We extend these investigations to include: (i) dry twin interface (two rotated single crystals in direct contact), (ii) crystal–water interfaces, (iii) crystal–protein–water systems. Our approach involves parametrization of interatomic potentials for multiple constituent phases and their interfaces, followed by systematic mechanical testing to extract relevant material properties. The following sections present our atomistic modelling.

### 2.1. Interface Testing Based on Cauchy Stress Vector

We use Cauchy’s theorem to characterise the interface. For an interface with normal vector **n**, the traction vector **t** acting on the interface is **t** = *σ ·* **n**. Here **t** comprises a normal stress (*σ*_*nn*_) and two orthogonal shear stresses (*τ*_*ns*_, *τ*_*nt*_), where *s* and *t* represent two orthogonal directions tangent to the interface plane (Figure 2(a)). **t** enables characterization of interfacial failure through systematic loading of interface systems in three modes and, consequently, constructing failure surface (Figure 2(b)). These modes include tensile loading perpendicular to the interface (*σ*_*nn*_ > 0, *τ*_*ns*_ = *τ*_*ns*_ = 0), shear loading parallel to the interface in two orthogonal directions (*σ*_*nn*_ = *τ*_*nt*_ = 0, *τ*_*ns*_ > 0 and *σ*_*nn*_ = *τ*_*ns*_ = 0, *τ*_*nt*_ > 0), as shown in Figure 2(c-e).

**Figure 2:**
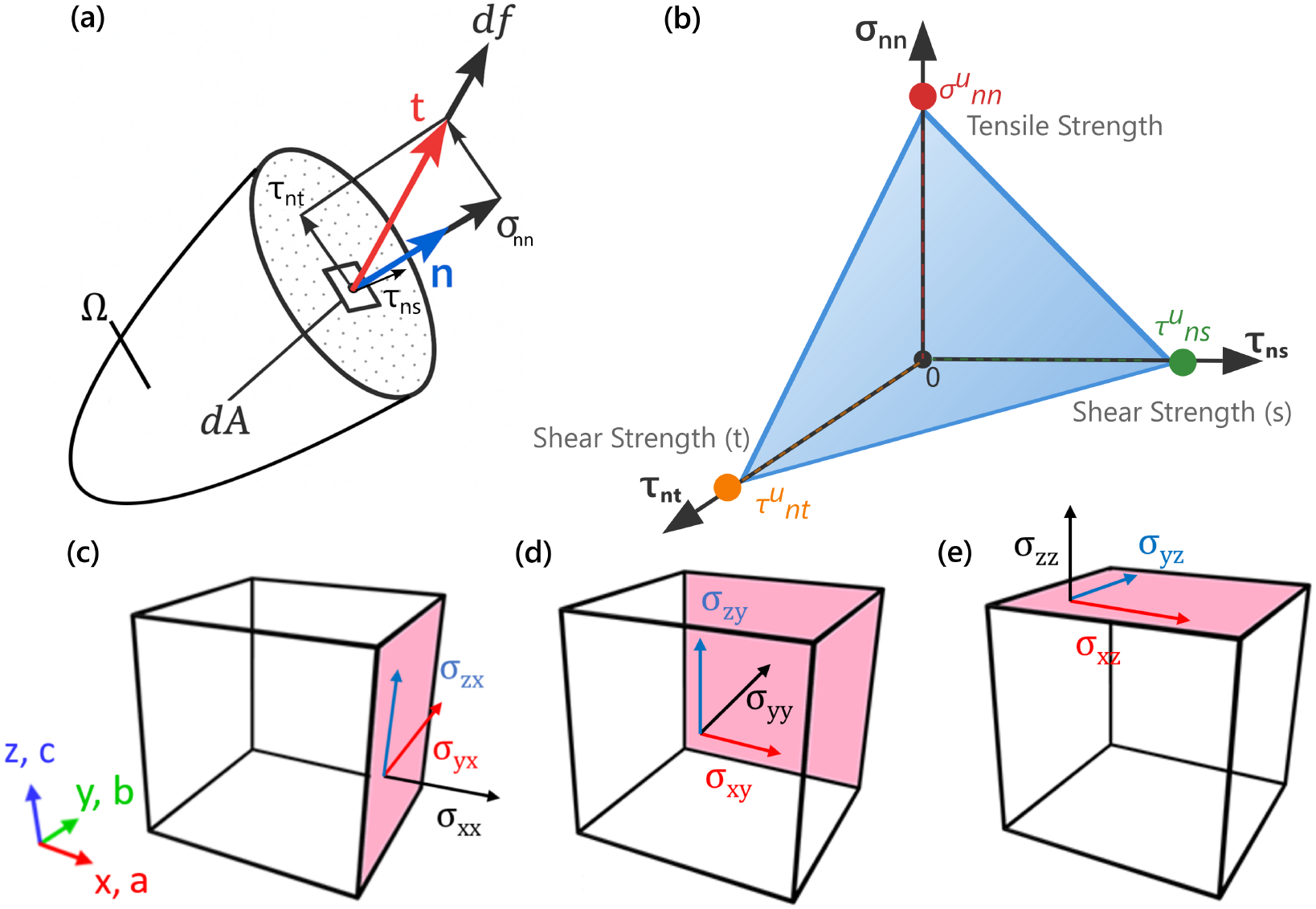
Interfacial contact between aragonite and solvated protein. (a) Decomposition of interface traction vector **t** into normal (*σ*_*nn*_) and two orthogonal shear components (*τ*_*ns*_, *τ*_*nt*_) following Cauchy’s theorem. (b) Three-dimensional failure surface showing triangular failure plane intersecting stress axes at pure tension strength 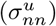 and pure shear strengths 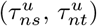. (c)-(e) Crystallographic interfacial planes: (c) {100} (*a*-axis normal), (d) {010} (*b*-axis normal), (e) {001} (*c*-axis normal), with corresponding stress vectors for each orientation

Employing a Mohr-Coulomb failure criterion for the interface (Equation (1)) [11], the failure surface forms a triangular plane in *σ*_*nn*_-*τ*_*ns*_-*τ*_*nt*_ space, intersecting the three axes at the pure tension strength 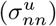, and the two shear strengths 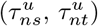. This failure criterion allows us to quantify interface failure under arbitrary stress states and to compare mechanical competence across different interface types, crystallographic orientations, and length scales.

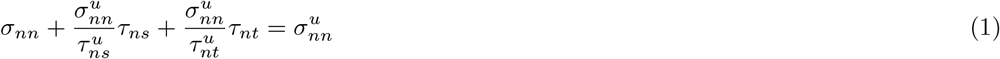

Here, 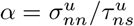 and 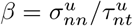 are the tensile-to-shear strength ratios, corresponding to the slopes of the two-dimensional Mohr–Coulomb criterion projections. Their inverses (1*/α* and 1*/β*) represent shear-to-tensile ratios, indicating shear load-bearing capacity relative to tensile strength. To quantify directional strength variation, we define a normalized strength anisotropy parameter:

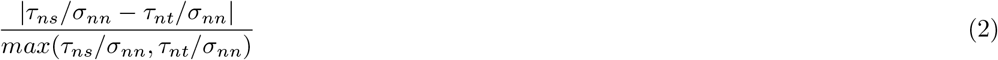

This parameter approaches zero when the two shear strengths are similar relative to tensile strength (indicating isotropic shear response), and increases toward unity as directional shear strength differences become more pronounced. It provides a dimensionless metric for comparing anisotropy across different interface types and orientations.

### 2.2. Interface Modelling and Simplification Rationale

Real coral interfaces exhibit complex three-dimensional morphologies with variable compositions, thickness distributions, and crystallographic relationships as can be seen from EBSD results [20, 37]. However, systematic investigation of interface mechanics requires well-controlled geometries that isolate specific physical mechanisms.

We investigate three limiting interface compositions that capture the range of possible behaviour: (i) dry twin boundaries; (ii) organic interfaces containing solvated CARPs; and (iii) water interfaces. The preparation of simulation systems containing solvated proteins in contact with aragonite crystal surfaces involves the following sequential steps. (i) We begin by generating an initial simulation box containing a solvated protein with `CHARMM`-integrated force field parameters [52] (Figure 3(a)). (ii) Steered Molecular Dynamics (SMD) is then applied to straighten and untangle the protein structure following the methodology of Zhang et al. [43], ensuring maximal exposure of functional groups for interaction with aragonite surfaces (Figure 3(b)). (iii) The system is subsequently processed with custom Python scripts to remove water molecules, rescale the simulation box, construct complex interface systems by implementing multiple protein copies at crystal interfaces, and position the interface in contact with an aragonite crystal surface (Figure 3(c-e)). (iv) Finally, the protein-crystal system is re-solvated using the TIP3P water model [53] within `LAMMPS`, followed by energy minimization, relaxation, and appropriate equilibration prior to mechanical testing.

**Figure 3:**
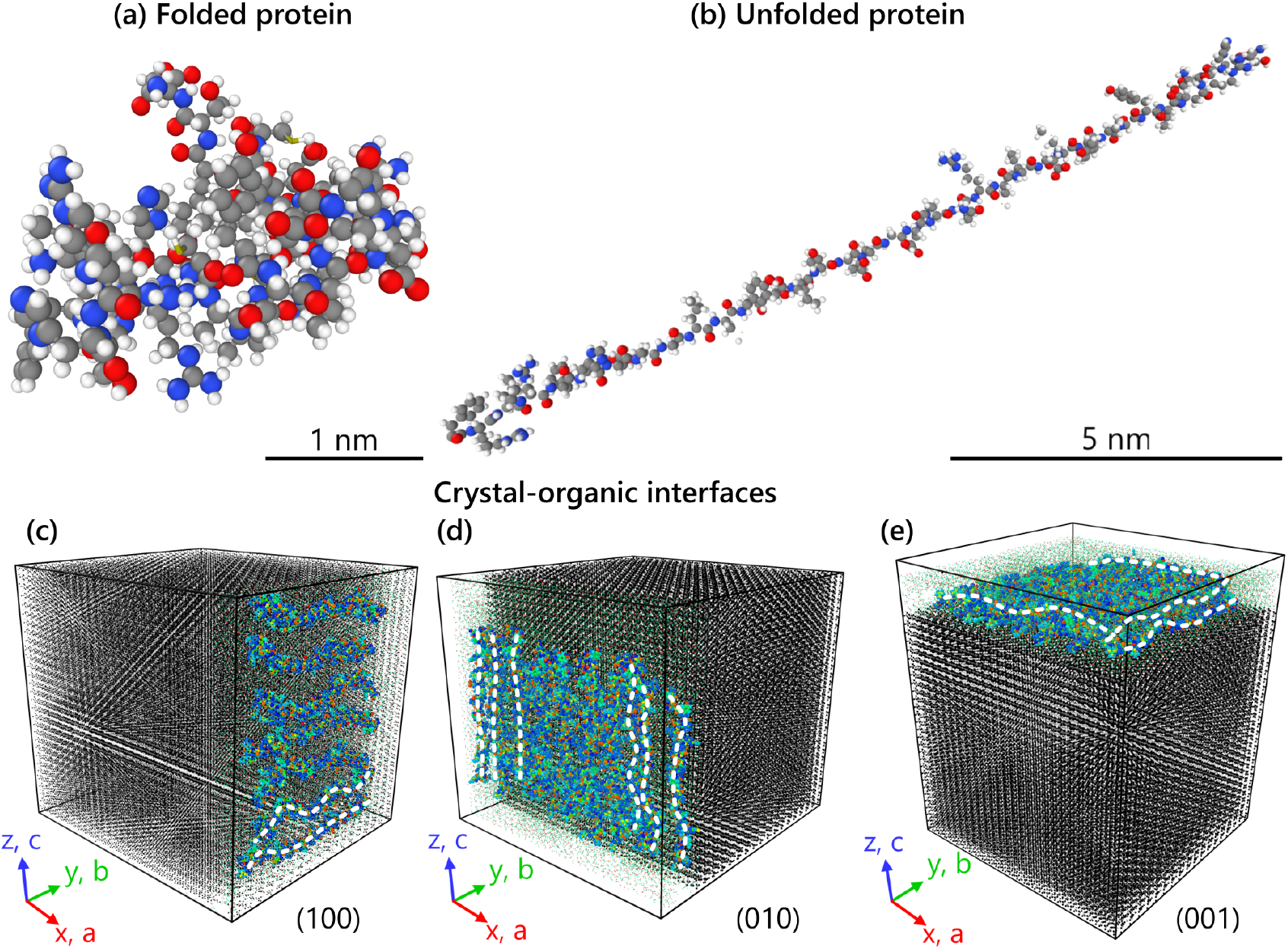
Interfacial contact between aragonite and relaxed solvated protein. (a) Initial folded protein. (b) Unfolded protein after applying SMD. Water molecules are not shown for clarity (c)-(e) Examples of an explicit interface between aragonite crystal and multiple unfolded proteins. Black atoms belong to aragonite crystal, coloured atoms attached to the surfaces are of the protein, smaller scattered atoms at the interface belong to water. Three configurations represent three types of considered interfaces according to Fig. 2(b-d) formulation: (c) single-layer interface at {100} (*x* −direction, *a* aragonite axis), (d) two-layer parallel interface at {010} (*y* −direction, *b* aragonite axis), and (e) two-layer perpendicular interface at {001} (*z* −direction, *c* aragonite axis). White dashed lines represent protein backbones

All relevant contact planes are assumed to be flat (Figure 3(c-e)). While our planar interface geometries cannot capture the full complexity of real CWC crystal interfaces, they represent limiting cases that enable isolation of fundamental mechanical mechanisms and provide quantitative bounds for interface properties. This approach agrees with established materials science methodologies where simplified geometries are used to understand complex systems [54, 55].

To explore the effect of crystallographic orientation on interface behaviour, we have constructed a set of simulation systems based on the aragonite unit cell structure. For each interface configuration (protein-crystal or water-crystal), we model three distinct orientations by attaching the interface along the {100}, {010}, {001}, and {110} crystallographic planes. Due to our coordinate system alignment, these correspond to interfaces perpendicular to the *x*−, *y*−, and *z*−directions of the simulation box, respectively, as illustrated in Figure 3(c-e). This approach allows us to quantify the anisotropic nature of interfacial interactions and establish structure-property relationships.

The dry twin configurations are constructed by rotating two single crystals by 28.16^°^(creating a misorientation of 56.32^°^) around the *c*-axis of the crystal (*z*-axis of the simulation box). These rotated segments are then joined to form a < 001 > {310} twin interface, with either direct contact or with protein/water at the boundary. A secondary twin configuration, < 001 > {110} with a misorientation angle of 63.82^°^, is also considered for comparative analysis.

### 2.3. Interatomic Potential Optimization

We use the Buckingham/Born model developed by Pavese et al. [36] as our starting point, as it has been validated for capturing both tensile and shear behaviour in aragonite through extensive comparison with experimental elastic constants and lattice dynamics calculations [36] (detailed potential formulations and parameters provided in Appendix A).

To ensure reproducibility and avoid arbitrary parameter adjustment, we employ a systematic optimization protocol based on experimental target properties. Our parameter refinement follows a three-step approach that begins with (i) target property selection based on experimentally established elastic constants*C*_*ij*_ (Table 1: Voigt [35], median values reviewed in Wolfram et al. [11]), aragonite lattice parameters (*a, b, c*), and unit cell volume [31]. We then perform (ii) systematic fitting using the General Utility Lattice Program (GULP) [56] with least-squares minimization through an iterative refinement procedure. Starting withPavese et al. [36] parameters as initial values, we first fit the diagonal stiffness components ((*C*_11_, *C*_22_, *C*_33_) as these are most reliably measured experimentally. The off-diagonal terms are subsequently refined while ensuring positive definiteness of the stiffness matrix, with continuous validation that lattice parameters remain within 1% of experimental values. Finally, (iii) the optimized parameters are validated against independent mechanical properties not used in the fitting process, ensuring transferability beyond the optimization dataset.

For the calculation of stiffness tensor we employ the standard Voigt-Mandel notation for the 6*×*6 stiffness matrix representation [57], where stress and strain vectors are defined as: 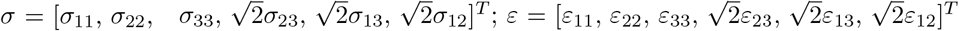. The elastic constants *C*_*ij*_ are extracted from three uniaxial tension tests (along *x, y, z* directions) and three shear deformation modes (*xy, xz, yz*), following the procedure established by Clavier et al. [58].

### 2.4. Proteins and Potentials

To model crystal-organic interfaces, we incorporate well-characterized aragonite-associated proteins from molluscan biomineralization systems. Specifically, we use structures of Lustrin A (Protein Data Bank (PDB): 1L3Q) [59] and Aragonite Protein-7 (PDB: 2JYP) [60]. These proteins were selected as representative models for crystal-organic interface studies because they are structurally well-characterized proteins known to mediate aragonite mineralization processes. Lustrin A (1L3Q) is a key structural protein from abalone nacre that serves as an established model for understanding protein-aragonite interactions [61, 62], while AP7 (2JYP) is specifically identified in coral skeletal formation and represents the direct coral protein-mineral interaction [63]. While these proteins originate from molluscan rather than coral systems, they provide validated structural models for investigating the fundamental mechanisms of protein-mediated aragonite formation at crystal-organic interfaces, which share common principles across different biomineralizing organisms.

At the atomic scale, the interactions between aragonite and different proteins are governed by similar interatomic forces (hydrogen bonding, electrostatic interactions, van der Waals forces) regardless of specific biological function, as all proteins consist of similar atomic constituents (C, N, O, H) with comparable functional groups (carboxyl, hydroxyl, amino). This approach enables systematic investigation of protein-mineral interface mechanics using established structural models while acknowledging that specific coral proteins may exhibit quantitative differences in binding strength and organization.

When modelling large molecules using MD, several options for force field description are available, all of which require additional intermediate steps in force field implementation and parameter adjustment. There are many degrees of freedom to control: parameters for non-bonded pair interactions, parameters for bonds, angles, dihedrals, impropers, etc. `CHARMM-GUI` [64] allows the conversion of protein structures directly from the PDB data format into `LAMMPS` native format, as well as preparing and converting force fields for protein interatomic interactions. The detailed implementation of protein force field is provided in Appendix B.

The proteins are relaxed and equilibrated in water, and prepared for integration with aragonite surfaces (Figure 3(a)). This computational approach enables systematic investigation of protein-mineral interactions at the molecular level. While the simplified model system cannot capture all aspects of the complex in vivo environment, it provides a controlled framework for examining fundamental interaction mechanisms between organic components and aragonite surfaces.

### 2.5. Simulation Setup

All Molecular Dynamics simulations were carried out using open-source software `LAMMPS` [65]. The simulations employed periodic boundary conditions in all directions, effectively modelling an infinite array of aragonite crystals and interfaces and enabling the study of bulk mechanical properties [66].

The aragonite crystal structure is characterized by an orthorhombic unit cell (Figure 1(a)) with lattice parameters *a* = 4.961*Å, b* = 7.967*Å*, and *c* = 5.740*Å*, containing 4 calcium, 4 carbon, and 12 oxygen atoms (20 atoms total per unit cell). In our simulation setup, the crystallographic axes are oriented with the *a*-axis along the *x*-direction, *b*-axis along the *y*-direction, and *c*-axis along the *z*-direction of the simulation coordinate system (Figure 1(a)). This orientation choice facilitates the systematic investigation of anisotropic mechanical properties across different crystallographic planes, as the three principal loading directions (*x, y, z*) correspond directly to the [100], [010], and [001] crystallographic directions, respectively.

Before performing mechanical tests, all systems were equilibrated using the Nosé-Hoover thermostat in the isothermal-isobaric ensemble (NPT) at 298 K and 1 bar for 48 ps. The time-step was set to 0.001 ps for all simulations. This equilibration protocol ensures proper relaxation of atomic positions and stress states before mechanical loading.

For tensile and shear tests, we used rectangular simulation boxes containing 35000 atoms for single-crystal tests and up to 250000 – 350000 atoms for dry twins and interfaces. The box dimensions ranged from approximately 60 *×* 97 *×* 70 *Å* for single-crystal systems up to 180 *×* 180 *×* 190 *Å* for twin and interface configurations, ensuring comparable aspect ratios and sufficient interface area for stress averaging. These system sizes provide adequate statistical sampling while maintaining computational efficiency. Simulation box loading was performed by applying an explicit deformation approach in which the dimensions of the simulation box were changed at a selected constant engineering strain rate (*ε*_*eng*_ = 10^10^*s*^−1^). Although these rates exceed experimental conditions due to fundamental computational constraints [67, 68], they provide quasi-static loading within computational limits. Strain was applied using the deform fix in `LAMMPS`, with engineering strain subsequently converted to true strain for analysis, according to *ε*_*true*_ = ln(1 + *ε*_*eng*_). Stress was computed using the standard virial formulation, which directly yields true stress in Molecular Dynamics simulations [69–71]. The strength of each system was defined as the maximum stress observed in the stress–strain response for each loading mode.

Atomistic structure analysis and visualization of results were performed using OVITO (Open Visualization Tool) [72]. This software enabled detailed examination of deformation mechanisms, displacement patterns, and structural changes during mechanical loading. Deformation analysis was performed by tracking atomic displacement patterns and local coordination environments throughout the loading process. Displacement magnitude calculations enable identification of deformation bands and phase boundaries, as well as changes in atomic arrangements indicative of phase transformations. The formation of deformation bands is characterized by regions of correlated atomic motion perpendicular to the loading direction, while phase transformations are identified through systematic changes in local atomic coordination that deviate from the original aragonite and protein structure (coordination numbers and bond angles).

To investigate the potential influence of atomic-scale defects on mechanical behaviour, we examined aragonite crystals with artificially introduced vacancy clusters (nanopores). While these idealized nanoscale defects do not directly represent the complex porosity observed in natural biominerals, they illustrate how atomic-scale discontinuities affect mechanical response. At the scale investigated (tens of nm), these defects represent vacancy clusters rather than true porosity and their mechanical effects may be limited compared to larger-scale structural features.

Two representative defect levels were investigated: 3% atomic vacancy concentration with 1 nm cluster diameter and 4% with 2 nm cluster diameter. Nanopores were introduced by randomly removing atoms within spherical regions while preserving overall electroneutrality. Larger pores (>2 nm) were found to be unstable due to long-range Coulombic distortions, whereas the chosen 1-2 nm sizes represent the maximum defect dimensions that maintain structural and electrostatic stability under equilibration.

### 2.6. Interface Architecture and Energetic Analysis

To systematically investigate the effects of protein arrangement and interface thickness on mechanical and energetic properties, we constructed multiple interface architectures. (i) Single-layer interfaces that feature protein molecules positioned in direct contact with the aragonite surface, creating thin interfacial regions with protein molecules in close proximity to the mineral surface (Figure 3(c)). (ii) Two-layer parallel interfaces consisting of two copies of the single-layer setup arranged in parallel alignment within thicker interfacial regions, creating increased protein density and greater protein-surface separation (Figure 3(d)). (iii) Alternatively, two-layer perpendicular interfaces orient two single-layer setups perpendicular to each other within the interface, providing alternative geometric arrangements while maintaining increased interface thickness (Figure 3(e)). Finally, (iv) mixed protein interfaces contain multiple protein types in a two-layer setup (viz., 2JYP+1L3Q) to investigate the effects of protein diversity on interface properties.

We primarily focus on single- and two-layer configurations as they are consistent with the observed ~1 nm interface thickness in coral aragonite [21], while also remaining computationally tractable, as increasing the number of protein layers significantly increases computational demands.

To address the fundamental questions about protein-mineral adhesion mechanisms, we quantify both structural and energetic aspects of interface behaviour. To quantify the role of water molecules in protein-mineral adhesion, we analyse: (i) identification of bridging water molecules that simultaneously coordinate with surface calcium ions (within 3.0 *Å*) and protein atoms (within 3.5 *Å*) and facilitate protein-mineral interactions, (ii) quantification of water utilization efficiency, defined as the percentage of interfacial water molecules that simultaneously bridge proteins and aragonite Ca atoms, as shown in Equation (3), and (iii) analysis of water density profiles and organisation patterns across different interface architectures.

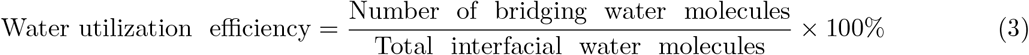

Interface adhesion strength is characterized through systematic binding energy analysis using direct binding energies quantify protein-mineral interactions:

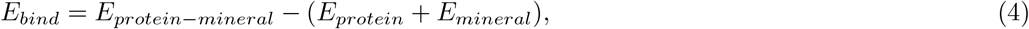

where *E*_*protein*−*mineral*_ represents the total energy of the combined system, and *E*_*protein*_ and *E*_*mineral*_ are the energies of isolated components in their equilibrium configurations. In addition, we calculate both area-normalized binding energies that accounts for interface size differences (Equation (5)) and per-residue binding efficiency that evaluates protein utilization (Equation (6)).

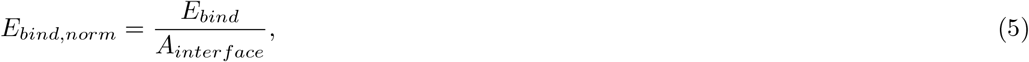

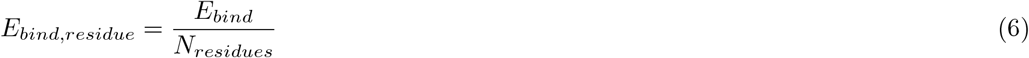

Here *A*_*interface*_ is the interface-mineral contact area, *N*_*residues*_ is the total number of amino acid residues in the interface.

Water-mediated interfacial energetics represent the comprehensive energetic stabilization including all protein-water, mineral-water, and protein-mineral interactions within the hydrated interface:

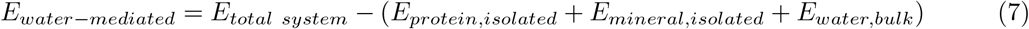

To understand how interfaces respond to mechanical stress, we track binding energy evolution during tensile loading, analysing: (i) initial binding optimization at low strains, where interfaces adjust to applied loads, (ii) progressive bond breaking and reorganization patterns, and (iii) the relationship between binding energy changes and mechanical stress-strain response. This analysis reveals whether interface failure occurs through sudden adhesion loss or gradual bond reorganization.

All interface analyses were performed across multiple crystallographic orientations to capture anisotropic effects, as shown in Figures 2(b-d) and 3(c). Interfaces were constructed along {100}, {010}, {001} crystallographic planes, corresponding to interfaces perpendicular to the *x*−, *y*−, and *z*−directions of the simulation box, respectively. Comparing direct binding energies, area-normalized values, and per-residue efficiencies across different surface orientations we identify crystallographic surfaces that provide optimal protein-mineral adhesion, thus, defining binding energy hierarchies.

This enables us to systematically compare of interface performance across different compositions (twin boundaries, protein interfaces, water interfaces), crystallographic orientations, and architectural arrangements, i.e., thickness of the interface, providing the quantitative foundation for developing structure-property relationships.

## 3. Results

The following sections establish how interface composition controls the transition from single-crystal strength (4-6 GPa) to bulk skeletal strength (~0.5 GPa). We begin with single-crystal aragonite properties, validating our refined force field and identifying orientation-dependent deformation mechanisms. We then characterize protein-mineral interface energetics, revealing how water mediates adhesion and how crystallographic orientation affects binding strength. Finally, we quantify interface mechanical response under tension and shear, constructing failure criteria that span nearly an order of magnitude in strength from twins (6.5 GPa) to ductile protein layers (0.6 GPa).

### 3.1. Single-Crystal Aragonite Properties

Our refined Pavese-based potential reproduces experimental elastic constants within 5% (Table 1), substantially improving upon the original parametrization [36]. For example, *C*_33_ component of the stiffness tensor is reproduced within 0.9% (experimental 85.0 GPa vs. simulated 84.2 GPa), whereas the original potential deviated by nearly 20% (68.3 GPa). This close agreement, detailed in Appendix C along with full mechanical properties, validates our approach for investigating interface mechanics.

Single-crystal aragonite exhibits anisotropic mechanical response [11], but the atomic-scale deformation mechanisms underlying this anisotropy remained unclear. Our simulations reveal that aragonite’s orthorhombic symmetry produces three distinct deformation pathways depending on loading direction. All orientations begin with elastic deformation uniform across all orientations (Figure 4(a,d,g)), but each crystallographic direction activates different mechanisms beyond the yield point.

**Figure 4:**
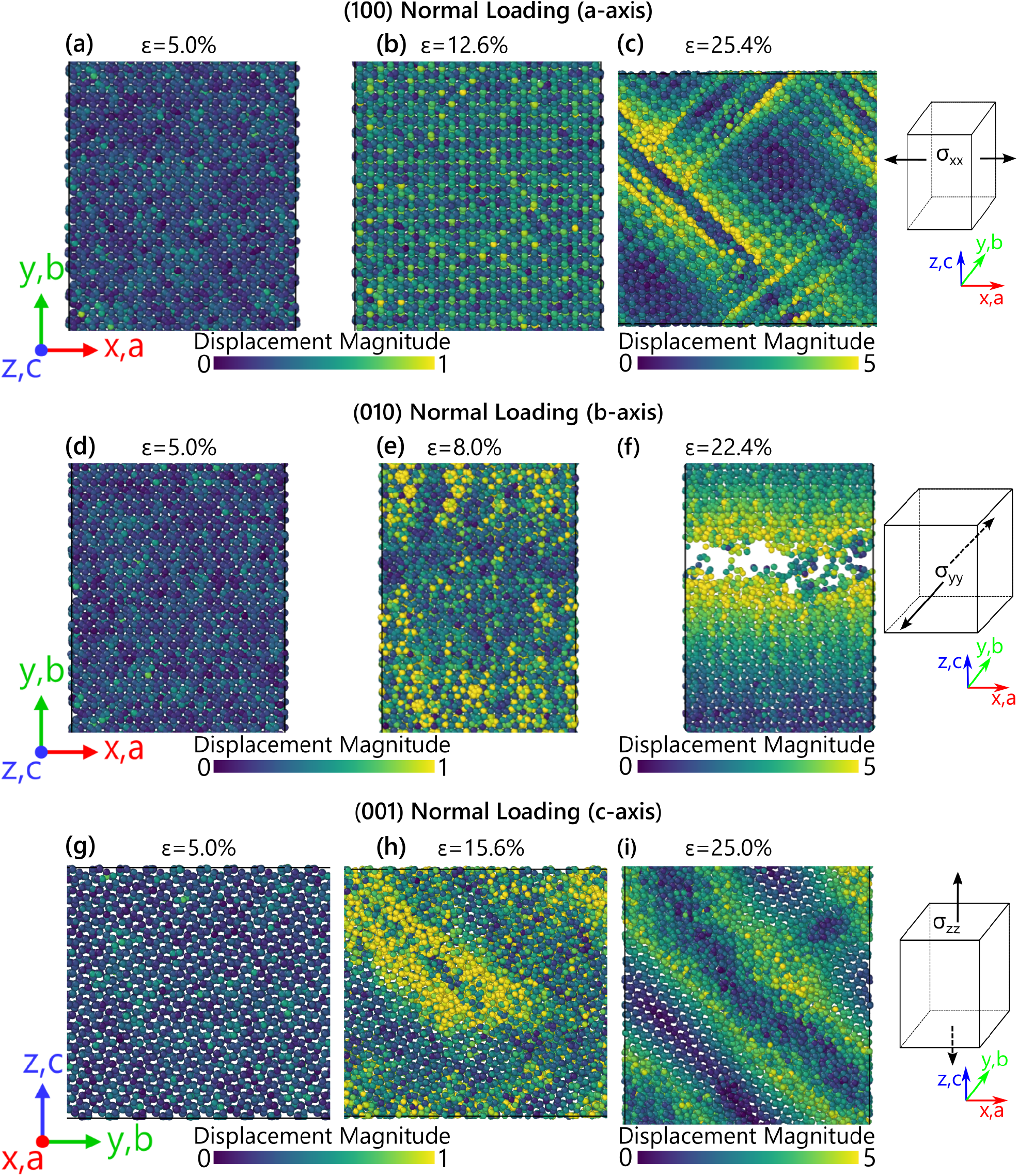
Orientation-dependent deformation mechanisms in single-crystal aragonite under tensile loading. Atomic structures coloured by displacement magnitude at three strain levels for each orientation. [100] loading (a-c): elastic deformation (5%), phase transformation (12.6%), and developed bands in transformed phase (25.4%). [010] loading (d-f): elastic deformation (5%), strain localization (8%), and crack propagation with voids (22.4%). [001] loading (g-i): elastic deformation (5%), band initiation following yield at 5.3 GPa (15.6%), and fully developed bands (25%)

Under [100] loading (along the *a*-axis), aragonite forms deformation bands at ~10% strain, then undergoes solid-state phase transformation at ~12% strain (Figure4(b)). By 25.4% strain, the transformed phase exhibits developed deformation bands (Figure 4(c)). Stress recovery accompanying this transformation demonstrates that the crystal accommodates large strains through atomic reorganization rather than fracture. In contrast, [010] loading (along the *b*-axis) produces brittle failure. After reaching peak stress (4.1 GPa), strain localizes (Figure 4(e)), rapidly evolving into crack nucleation and propagation with clear void formation by 22.4% strain (Figure 4(f)), offering no alternative deformation pathway. Along the [001] direction (*c*-axis, the coral growth direction), our simulations show deformation band initiation following yield at 5.3 GPa (Figure 4(h)). These bands develop fully with increasing deformation (Figure 4(i)), involving coordinated atomic rearrangements that maintain partial load-bearing capacity, producing the stress plateau observed in Figure C.1.

Shear loading produces similar deformation band formation (Figure 4(c,h,i)), confirming that aragonite accommodates shear through coordinated atomic reorganization rather than brittle fracture. Together, these results reveal three orientation-dependent mechanisms: deformation banding in [100] and [001] directions and shear deformation, brittle fracture along [010], and phase transformation specific to [100] loading.

Introducing nanopores produces contrasting effects on tensile versus shear strength (Table 2). Tensile strength increases relative to perfect crystals, most dramatically in the [010] direction (4.10 → 6.33, 6.39 GPa), while shear strength decreases slightly across all orientations. Vacancy clusters impede crack propagation in tension, requiring higher stresses to initiate failure, but promote shear localization, thereby reducing shear strength. This contrast demonstrates how the same microstructural feature affects strength differently depending on the active deformation mechanism.

**Table 2:** Aragonite strength for perfect crystal and crystals with nanoporosity in GPa. Values in parentheses show percentage change relative to the perfect crystal from this study

| System | $\sigma_{xx}$ | $\sigma_{yy}$ | $\sigma_{zz}$ | $\sigma_{xy}$ | $\sigma_{xz}$ | $\sigma_{yz}$ |
| --- | --- | --- | --- | --- | --- | --- |
| Perfect crystal<br>this study | 4.98 | 4.10 | 5.34 | 4.51 | 5.08 | 5.06 |
| Perfect crystal<br>from [11] | 4.97 | 4.00 | 5.33 | 3.92 | 4.06 | 4.03 |
| 3% porosity,<br>1 nm pores | 5.26<br>(+5.6%) | 6.33<br>(+54.4%) | 5.57<br>(+4.3%) | 4.26<br>(-5.5%) | 4.06<br>(-20.1%) | 4.26<br>(-15.8%) |
| 4% porosity,<br>2 nm pores | 5.52<br>(+10.8%) | 6.39<br>(+55.9%) | 5.62<br>(+5.2%) | 4.41<br>(-2.2%) | 4.32<br>(-15.0%) | 4.51<br>(-10.9%) |

Single-crystal aragonite maintains high strength even with nanopore defects, consistent with experimental measurements [11]. This strength exceeds bulk coral skeleton strength by an order of magnitude, confirming that crystal defects cannot explain the transition from 4-6 GPa single crystal strength to ~0.5 GPa polycrystalline strength. Instead, the interfaces linking aragonite grains must control macroscopic mechanical response. The following sections therefore characterize these critical interfaces: twins, protein interfaces, and water boundaries.

### 3.2. Interfacial Structure and Adhesion Mechanisms

This section characterizes protein-aragonite interface structure and adhesion energetics in fully equilibrated, stress-free configurations. We examine water-mediated binding mechanisms, crystallographic orientation effects, and interface thickness dependencies. These equilibrium properties provide the basis for understanding mechanical response under load (Section 3.3).

Protein-aragonite interfaces across all simulated configurations exhibit water-mediated binding as the dominant adhesion mechanism. A representative (010) interface (Figure 1(b)) shows proteins adhering to mineral surfaces through structured water networks that bridge protein functional groups and surface calcium ions. Water molecules organize into layers extending several *Å* from the mineral surface (blue dashed line in Figure 1(b)), creating high-density zones at 1.9-2.3 times bulk water density. Proteins maintain their hydration shells while establishing contact through extensive hydrogen-bonded networks. This water-mediated mechanism operates consistently across all simulated crystallographic orientations and protein arrangements.

Interface architecture strongly affects water bridging efficiency (Figure 5). Single-layer protein interfaces support 1.6-2.3 times more bridging water molecules than two-layer systems, achieving substantially higher utilization efficiency (8.9-17.2% vs. 2.1-3.0%; Equation (3)). This difference arises because water molecules in thicker interfaces must span larger protein-surface distances, reducing the probability of simultaneous coordination with both protein and mineral. Consequently, thicker interfaces sacrifice bridging efficiency for other structural benefits explored below.

**Figure 5:**
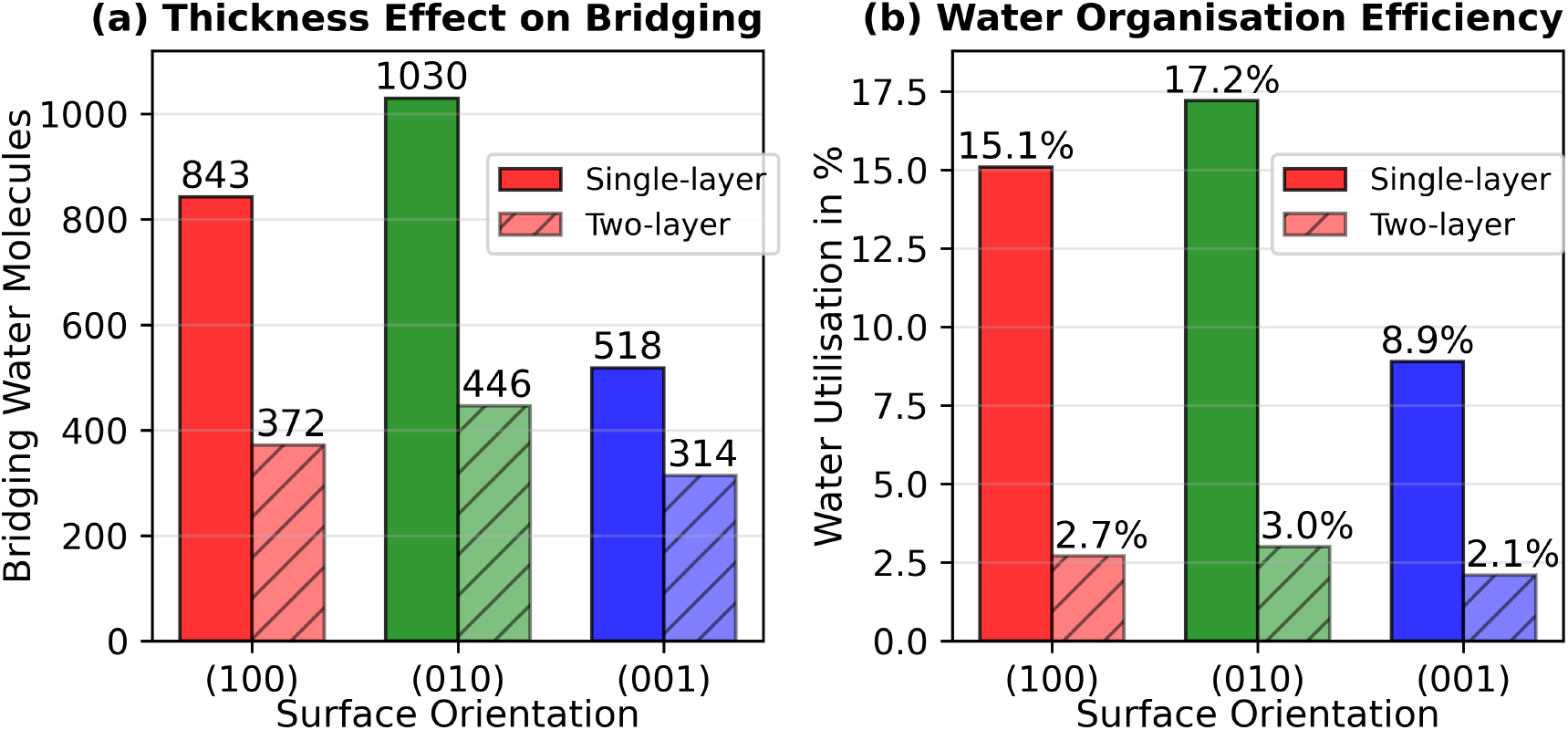
Interface architecture effects on water-mediated binding. (a) Comparison of bridging water molecules between single-layer (solid bars) and two-layer (hatched bars) protein interfaces across different surface orientations. (b) Water utilization efficiency showing the fraction of interfacial water participating in protein-mineral bridging

Crystallographic orientation also affects water utilization efficiency in our models. Single-layer interfaces exhibit the hierarchy (010) > (100) > (001), which correlates inversely with surface calcium density. The (010) surface achieves highest utilization (17.2%) by providing optimal spacing for water-mediated bridging between protein functional groups and dispersed calcium sites. In contrast, the (001) surface with its dense calcium packing shows only 8.9% utilization, as fewer water molecules can simultaneously coordinate both components. Two-layer systems maintain the same orientation hierarchy but with uniformly lower efficiencies due to geometric constraints discussed above.

Water utilization quantifies geometric bridging efficiency, but binding energies reveal total thermodynamic adhesion strength. These complementary metrics show that efficient water bridging does not necessarily produce the strongest adhesion, as direct electrostatic interactions also contribute significantly.

Binding energy calculations across different architectures and orientations (Table 3) reveal a clear hierarchy: (100) > (001) > (010). The (100) surface exhibits strongest adhesion in single-layer configurations (−2259 kcal/mol), followed by (001) surfaces (−1484 kcal/mol) and (010) surfaces (−1031 kcal/mol). This hierarchy reflects differences in calcium site density and electrostatic screening within our force field representation of protein-mineral interactions.

**Table 3:** Protein-aragonite binding energies across different interface configurations. Direct binding energies quantify protein-mineral adhesion strength, area-normalized values account for interface size differences, per-residue values indicate protein utilization efficiency, and water-mediated energies reflect total system reorganization in aqueous environments. Parallel (∥) and perpendicular (⊥) designations refer to protein alignment within two-layer interfaces. More negative values indicate stronger binding

| Surface | Interface Type<br>(# layers,<br>protein type) | Direct<br>Binding,<br>kcal/mol | Area-Normalized,<br>kcal/mol/$\text{\AA}^2$ | Per-Residue,<br>kcal/mol/res | Water-Mediated,<br>kcal/mol |
| --- | --- | --- | --- | --- | --- |
| (100) | Single, 2JYP | -2258.78 | -0.103 | -8.96 | -88780.26 |
|  | Single, 1L3Q | -552.16 | -0.140 | -11.50 | -15782.38 |
| | Two $\parallel$, 2JYP | -1974.05 | -0.084 | -3.92 | -115786.16 |
| | Two $\perp$, 2JYP | -1433.70 | -0.065 | -2.84 | -106572.56 |
|  | Two, 2JYP+1L3Q | -728.47 | -0.039 | -1.79 | -90219.24 |
| (010) | Single, 2JYP | -1031.44 | -0.045 | -4.09 | -66599.50 |
| | Two $\parallel$, 2JYP | -679.58 | -0.028 | -1.35 | -88295.25 |
| (001) | Single, 2JYP | -1484.25 | -0.068 | -5.89 | -58991.67 |
| | Two $\parallel$, 2JYP | -1025.31 | -0.044 | -2.03 | -81861.64 |

This binding energy hierarchy inverts the water utilization hierarchy established above. Water utilization follows (010) > (100) > (001), reflecting geometric efficiency of water-mediated bridging, whereas binding energy follows (100) > (001) > (010), reflecting total thermodynamic stability from both direct electrostatic and water-mediated contributions. This inversion demonstrates that surfaces optimizing water structure do not necessarily maximize total adhesion—direct protein-mineral interactions dominate overall binding strength.

Interface architecture also affects binding strength. Single-layer configurations consistently exhibit stronger binding than two-layer arrangements: the (100) surface shows 13% reduction when adding a second layer, while the (001) surface shows 31% reduction. Despite these thickness effects, area-normalized binding energies confirm that the orientation hierarchy (100) > (001) > (010) persists across all architectures. This consistency indicates that within our models, crystallographic orientation controls inherent protein-mineral affinity independent of interface thickness.

Per-residue binding efficiency reveals how effectively proteins utilize their amino acids for adhesion. Single-layer interfaces achieve highest efficiency (−8.96 kcal/mol per residue for (100) surfaces), while two-layer interfaces show reduced efficiency: −3.92 kcal/mol for parallel arrangements (∥) and −2.84 kcal/mol for perpendicular arrangements (⊥). This reduction reflects increased reliance on water-bridged adhesion in thicker interfaces. Comparing protein types, 1L3Q achieves higher per-residue efficiency than 2JYP despite lower total binding energy (−552 vs. −2259 kcal/mol), because its chains smaller size and higher active site density optimize surface contact.

These results indicate that within our simulated systems, interface performance depends on balancing two factors: geometric water bridging efficiency and total electrostatic binding strength. The (010) surface maximizes the former but shows weakest overall adhesion, while the (100) surface shows strongest adhesion through direct protein-mineral electrostatics despite intermediate water utilization. Interface thickness introduces an additional trade-off: single layers maximize binding per residue, while multiple layers distribute adhesion across more protein-mineral contacts at the cost of reduced per-residue efficiency. Section 3.3 examines how these equilibrium properties translate into mechanical performance under load.

### 3.3. Mechanical Response of Interfaces

Section 3.2 characterized interface adhesion energetics in equilibrium. This section examines mechanical response under tensile and shear loading. Because the (100) surface showed strongest binding, we performed comprehensive testing on all interface types at this orientation. We also tested representative configurations at (010) and (001) orientations for comparison, including single-layer 2JYP proteins, two-layer parallel 2JYP proteins, dry twin boundaries, and water interfaces (full results on (100) surface in Appendix D).

Figure 6 compares stress-strain curves of different interface types under tensile (normal) loading. Distinct mechanical responses emerge depending on both interface type and crystallographic orientation, consistent with the structural and energetic patterns observed in Section 3.2.

**Figure 6:**
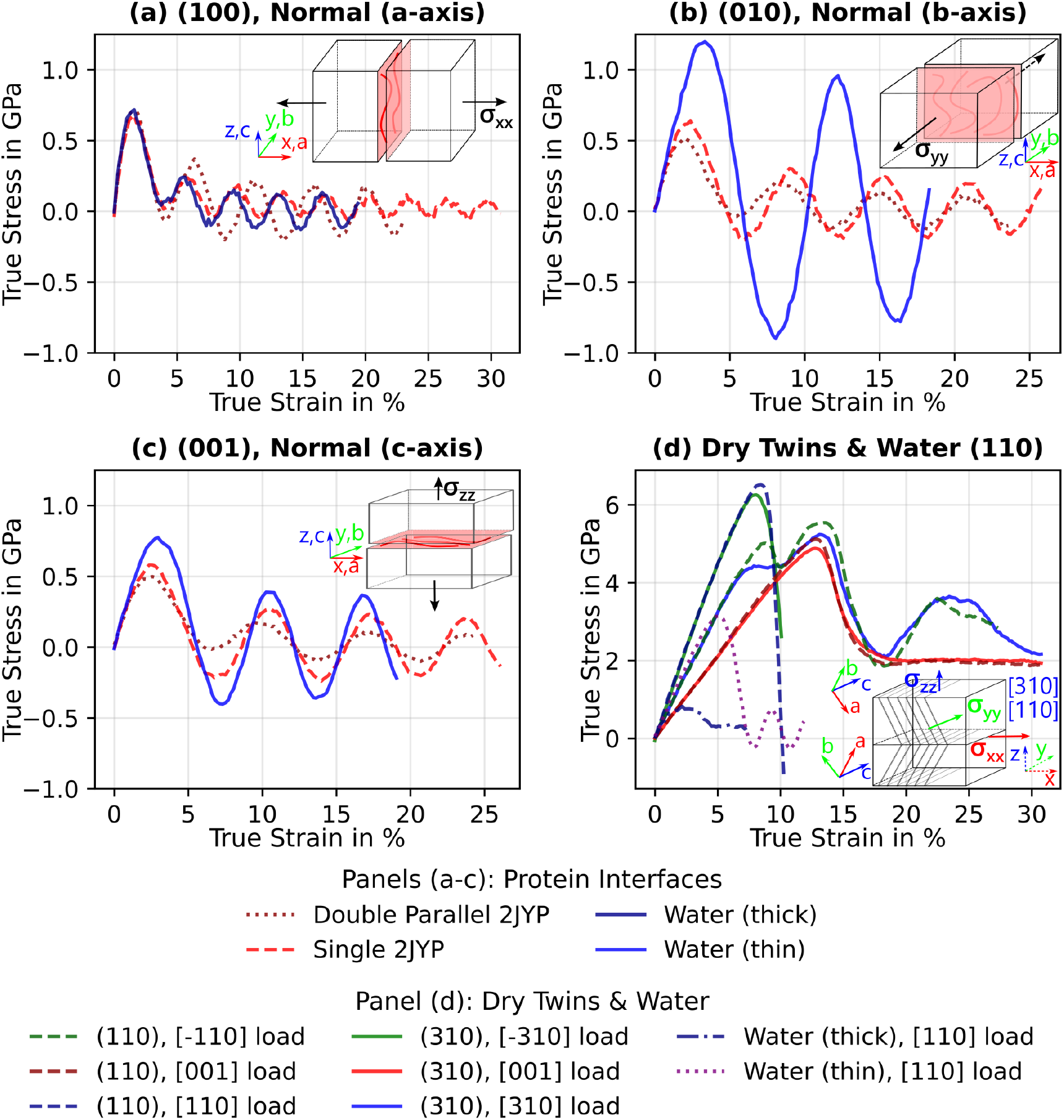
Tensile response of aragonite-based interfaces across crystallographic orientations. Loading directions correspond to aragonite crystallographic axes with interface orientations denoted by Miller indices (*hkl*): *x* ∥*a, y* ∥*b, z*∥ *c*. Insets show loading geometry for each orientation. (a) Simulation box *x*-direction loading on (100) interfaces showing protein and water responses, (b) *y*-direction loading on (010) interfaces, (c) *z*-direction loading on (001) interfaces, and (d) comparison of dry twin boundaries and water (110) interfaces across all loading directions. For twin and water (110) rotated systems [001] axis is the tilt axis for the interface

Loading along the *x*-direction (crystal *a*-axis, [100]) produces the highest protein-mediated interface strengths: 0.56-0.72 GPa (Figure 6(a)). Two-layer parallel 2JYP configurations reach 0.72 GPa, while single-layer interfaces achieve 0.56-0.68 GPa. These values align with the strong binding energies measured for (100) surfaces in Section 3.2, and the enhanced performance of two-layer systems may reflect cooperative hydrogen bonding between layers.

Loading in the *y*-direction (crystal *b*-axis, [010]) produces intermediate protein interface strengths (Figure 6(b)). Two-layer parallel 2JYP configurations reach 0.52 GPa, while single-layer interfaces achieve 0.64 GPa. Notably, pure water interfaces show the highest strength among all (010) configurations (1.2 GPa), consistent with the efficient water bridging observed for this orientation in Section 3.2.

Loading along the *z*-direction (crystal *c*-axis, [001], the coral growth direction) produces the lowest overall interface strengths (Figure 6(c)). Protein interfaces reach 0.49-0.58 GPa, consistent with the intermediate binding energies measured for (001) surfaces, while water interfaces reach ~0.78 GPa. The similar performance of single- and two-layer protein configurations suggests that interface architecture affects strength less strongly in this orientation, where surface properties may limit adhesion.

Twin boundaries exhibit markedly higher tensile strength, reaching up to 6.5 GPa, which is an order of magnitude greater than protein interfaces (Figure 6(d)). The (110) twins (63^°^misorientation) show slightly higher strengths (4.4-6.5 GPa) than (310) twins (56^*°*^) (4.9-6.3 GPa), potentially reflecting subtle differences in crystallographic matching across the boundary.

Water (110) interfaces display thickness-dependent strength: thin interfaces (< 5*Å*) reach >3.1 GPa, likely due to residual Coulombic attraction between mineral surfaces across the narrow water layer, whereas thick interfaces (> 10*Å*) exhibit ~0.8 GPa as water screening becomes dominant. Despite their lower strength, both configurations deform plastically and accommodate large strains, potentially functioning as compliant elements in skeletal hierarchies.

Figure 7 shows shear stress-strain responses of representative interfaces. Shear loading produces deformation mechanisms distinct from tensile loading, with lower absolute strengths and higher sensitivity to interface thickness.

**Figure 7:**
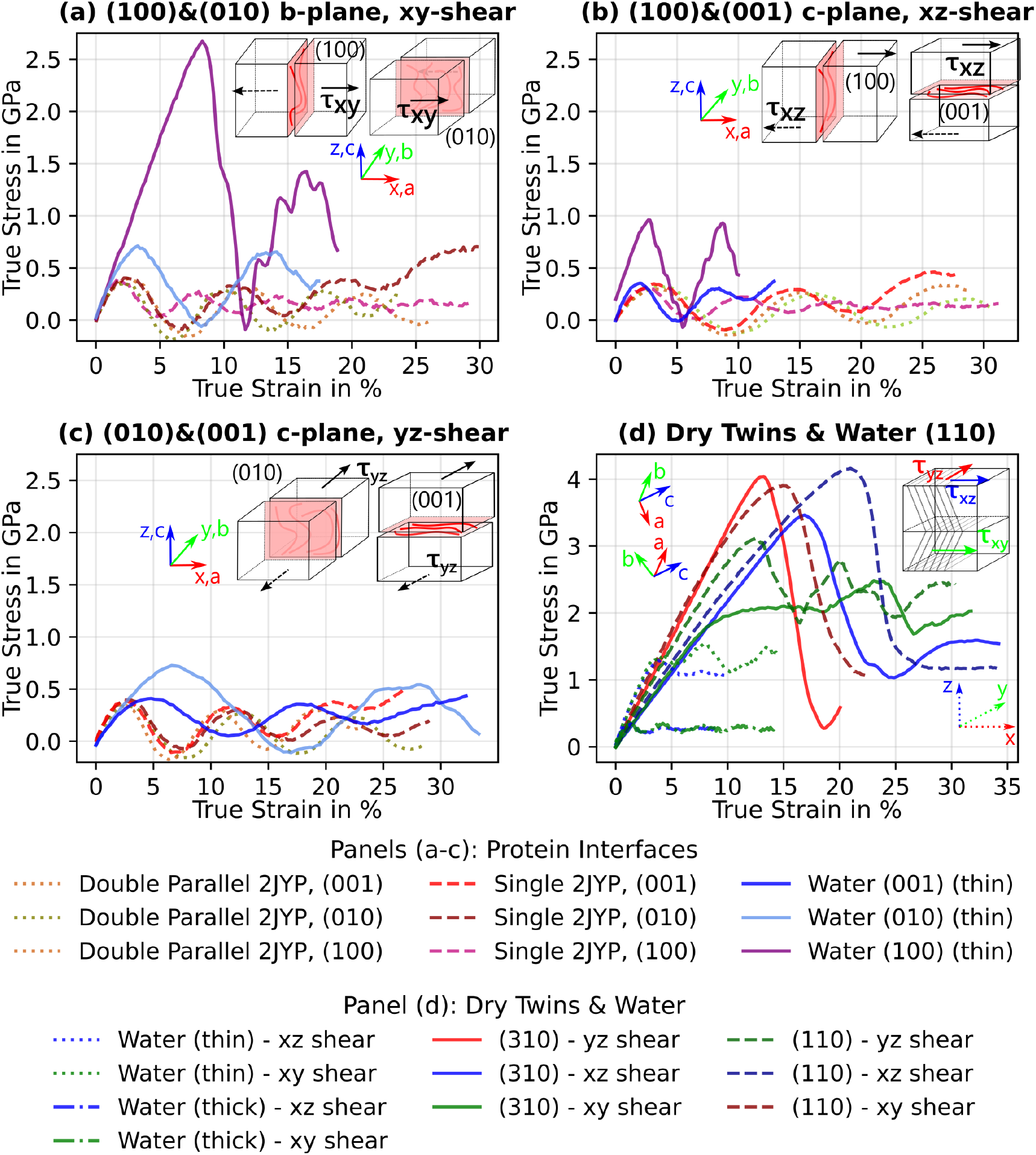
Shear response of aragonite interfaces across loading modes. Insets show shear loading geometry for each mode. (a), (b), (c) *xy* (crystal *b*-plane), *xz* (crystal *c*-plane), and *yz* (crystal *c*-plane) shear deformation modes showing representative protein and water interface behaviour. (d) Comparison of dry twin boundaries and water interfaces across all shear directions. For twin and water (110) rotated systems [001] axis is the tilt axis for the interface

Protein interfaces reach peak shear stresses of approximately 0.4 GPa for *xy* and *yz* shear, and 0.3 GPa for *xz* shear (Figures 7(a–c)). Single-layer 2JYP interfaces display orientation-dependent post-peak behaviour: (010) and (001) interfaces show stress recovery at higher strains, while (100) interfaces exhibit gradual deterioration of shear stresses.

Dry twin systems show that mechanical response during shear is governed primarily by deformation of the rotated crystal lattice, mirroring single-crystal behaviour, rather than interface-specific sliding (Figure 7(d)). Strength differences between (110) and (310) twins appear to reflect distinct lattice matching rather than interfacial bonding differences.

Water interfaces again show pronounced thickness dependence. Thin (110) water layers reach 1.3-1.4 GPa, whereas thick layers exhibit only 0.33-0.34 GPa. Across all orientations, thinner water layers appear to maintain stronger coupling between opposing crystal surfaces, while thicker layers facilitate sliding and energy dissipation.

Analysis of failure mechanisms shows that protein interfaces undergo progressive degradation through sequential bond breaking, evidenced by multiple stress peaks and gradual strength reduction that is more pronounced in tensile loading. This mechanism provides substantial energy dissipation. Water interfaces demonstrate controlled sliding with stress levels that vary significantly based on thickness, consistent with their potential role in accommodating large deformations while maintaining interface integrity. The thickness-dependent mechanical response supports the geometric effects on water bridging observed in Section 3.2.

To connect interface energetics and mechanical response, we explore the evolution of binding energy during tensile loading shown in Figure 8. The binding energy evolution exhibits a characteristic pattern that differs significantly from the mechanical stress-strain response. Binding energy tracks cumulative bond degradation smoothly, while stress reflects instantaneous load-bearing capacity and shows oscillatory behaviour as local protein-mineral contacts fail and rearrange.

**Figure 8:**
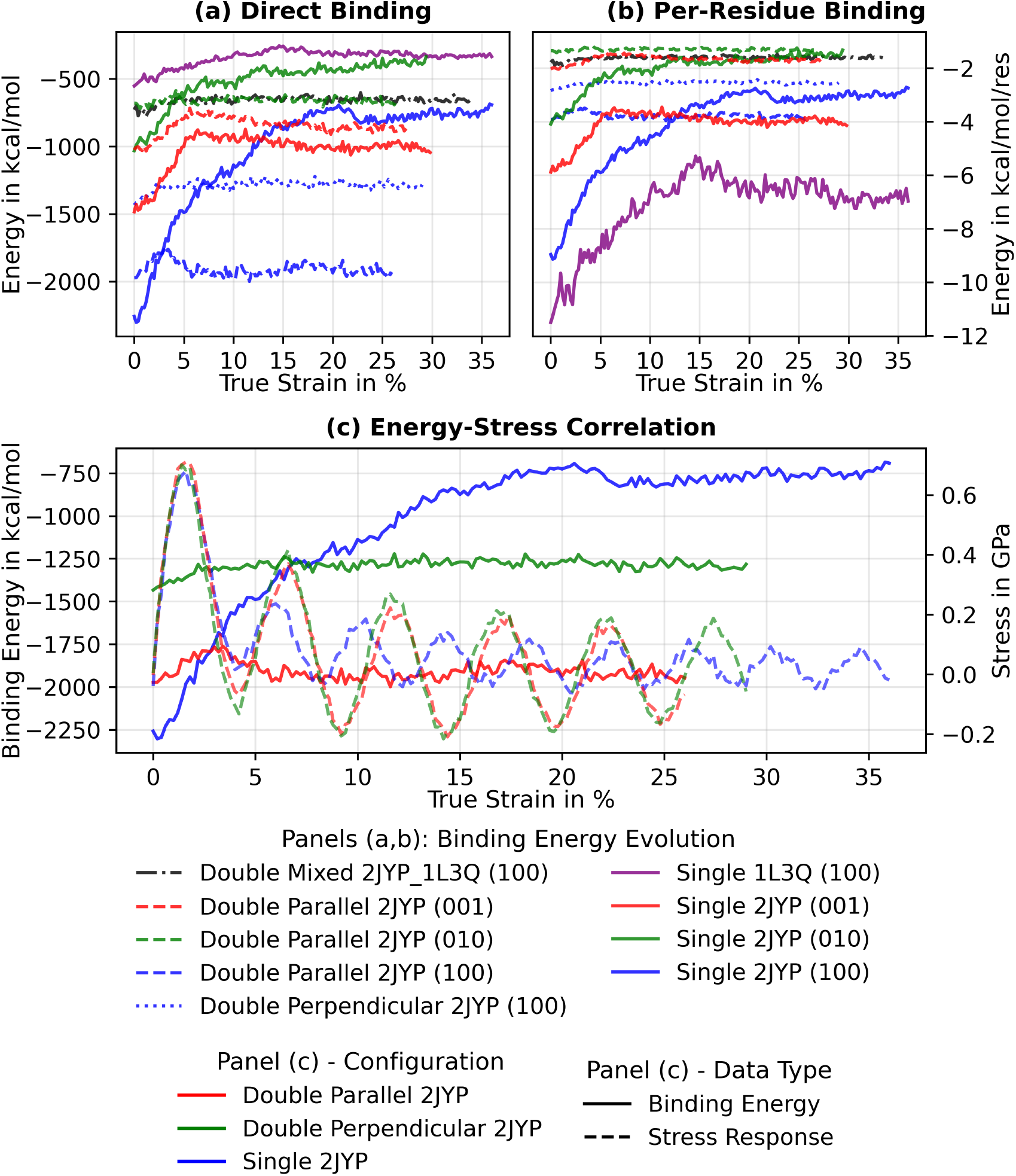
Binding energy evolution during tensile loading of protein interfaces. (a) Direct binding energy evolution shows progressive reduction. Single-layer interfaces (solid lines) exhibit more dramatic changes than two-layer configurations (dashed/dotted lines). (b) Per-residue binding efficiency. (c) Energy-stress correlation for representative cases shows smooth binding energy evolution compared to oscillatory mechanical response. Colours represent protein configurations: blue (single-layer ∥ 2JYP), red (two-layer ∥ 2JYP), green (two-layer ⊥ 2JYP). Line styles distinguish data types: solid (binding energy), dashed (stress response)

Binding energies decrease steadily with applied deformation but do not approach zero even beyond 20-30% strain, indicating that partial adhesion persists. This behaviour suggests that water-mediated hydrogen-bond networks reorganize rather than fail catastrophically.

Single-layer interfaces show the strongest initial adhesion but also the steepest weakening, consistent with greater sensitivity to deformation in our models. Two-layer configurations exhibit more gradual energy decay and correspondingly smoother stress-strain responses, suggesting enhanced ductility through distributed adhesion.

Per-residue binding efficiency (Figure 8(b)) follows similar trends to total binding energy but with enhanced sensitivity to configuration differences. The correlation between energy and stress (Figure 8(c)) shows that energy loss proceeds more smoothly than stress fluctuations, indicating that interface failure in our models occurs through progressive bond rearrangement rather than abrupt bond breakage.

### 3.4. Quantitative Failure Analysis

The previous sections described interface mechanical response under tension and shear. This section quantifies relative performance across all interface types and constructs three-dimensional failure criteria using a Mohr-Coulomb approach (Section 2.1). This analysis allows direct comparison of load-transfer capability and mechanical anisotropy across organic, aqueous, and crystalline interfaces.

Comparative analysis of tensile and shear data from Section 3.3 (Figure 9) reveals a clear strength hierarchy spanning approximately one order of magnitude: from ~0.5 GPa (protein interfaces) to 6.5 GPa (dry twins). Dry twin boundaries rank strongest, followed by thin water interfaces, with thick water and protein interfaces showing similar but lower strengths.

**Figure 9:**
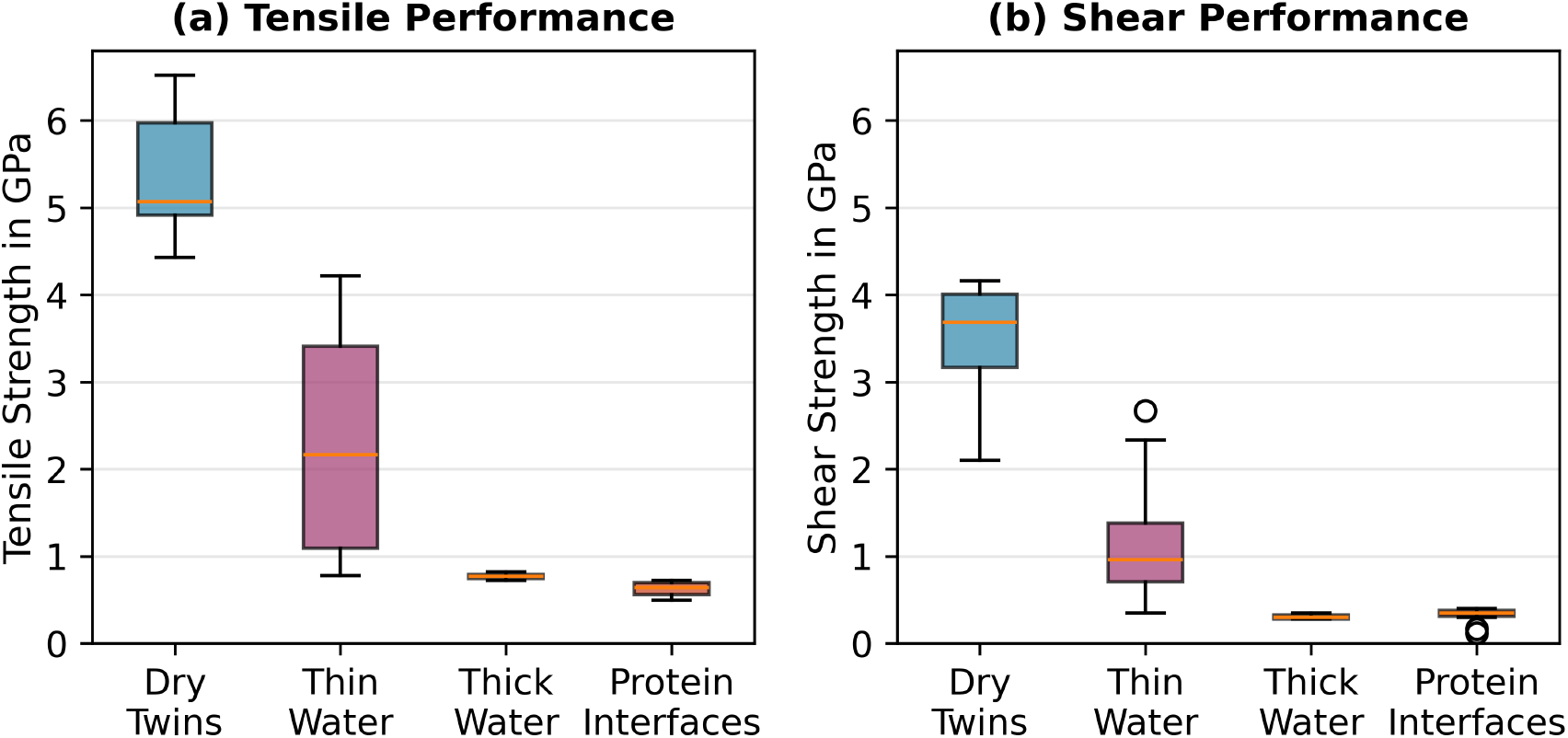
Tensile (a) and shear (b) strength comparison across interface types, spanning one order of magnitude in performance. Dry twin boundaries provide highest strength through direct mineral-mineral bonding, while water interface performance depends critically on thickness due to Coulombic screening effects. Bars indicate maximum and minimum values across multiple configurations.

This hierarchy correlates with interfacial bonding character. Twin boundaries involve direct ionic and covalent bonding between contiguous crystal lattices, producing maximal strength but lower ductility. Thin water interfaces maintain partial electrostatic coupling between aragonite surfaces, yielding relatively high strength with limited energy dissipation. Thick water and protein interfaces are dominated by hydrogen-bonded and screened electrostatic interactions, producing lower peak strength but enhanced ductility and energy absorption capacity.

Overall, interface performance reflects a trade-off between strength and deformation tolerance: stronger direct bonding produces more brittle response, while hydrogen-bonded and hydrated systems facilitate progressive, damage-tolerant deformation.

Tensile and shear data across three orthogonal directions enable construction of complete three-dimensional Mohr-Coulomb failure surfaces (Figures 10, 11). We quantify interface mechanics using shear-to-tensile strength ratios (*τ*_*ns*_*/σ*_*nn*_ = 1*/α, τ*_*nt*_*/σ*_*nn*_ = 1*/β* from Equation (1)) and the normalized strength anisotropy parameter (Equation (2)). These metrics characterize relative shear versus tensile load-bearing capacity and directional strength variation, respectively.

**Figure 10:**
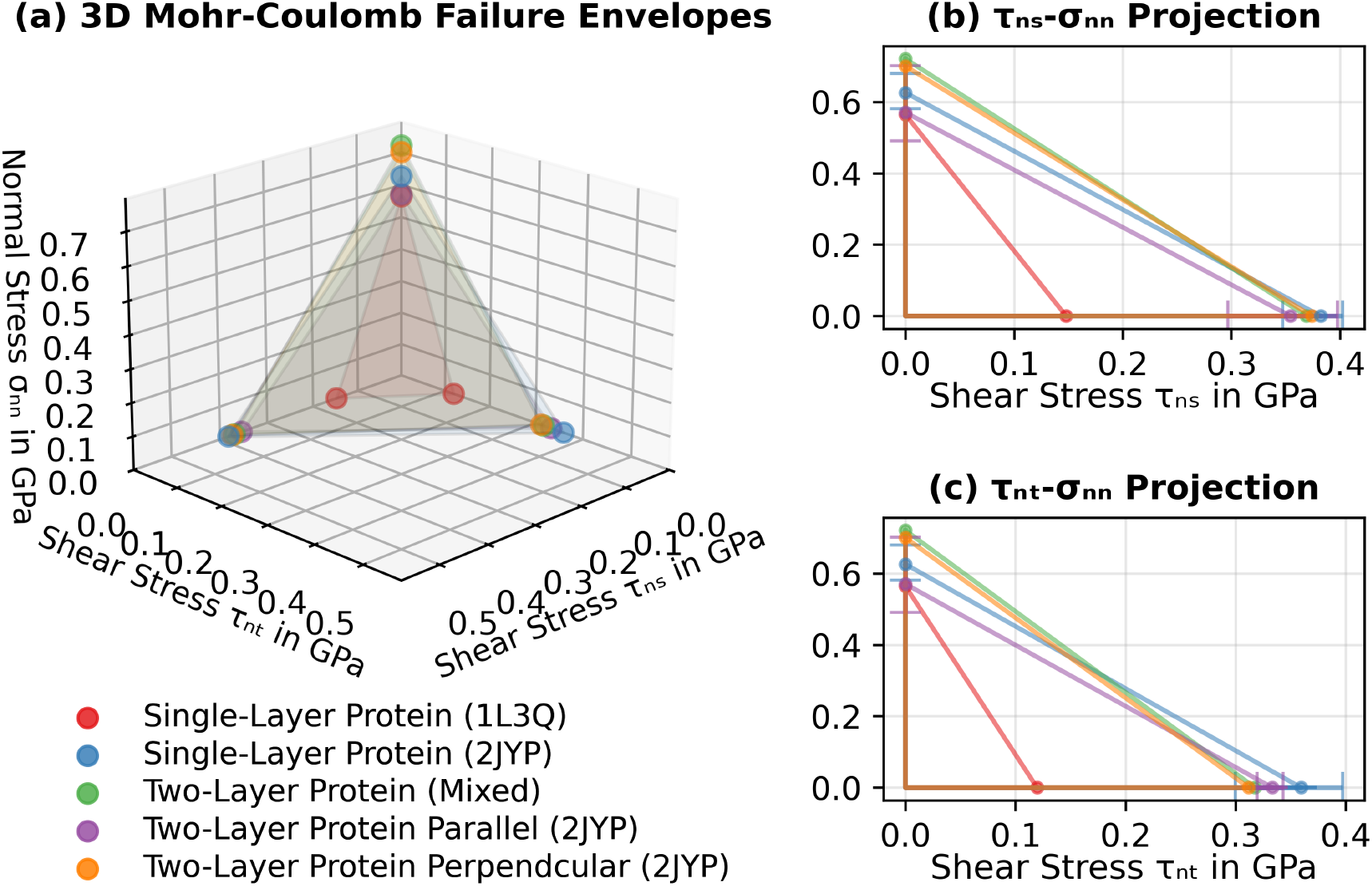
Three-dimensional Mohr–Coulomb failure surface for protein-mediated interfaces. (a) Full failure surface in *σ*_*nn*_-*τ*_*ns*_-*τ*_*nt*_ space, showing critical stress combinations leading to interface failure. (b), (c) Two-dimensional projections highlighting the relationships between normal and shear stress components. Bars indicate minimum and maximum strengths across multiple configurations

**Figure 11:**
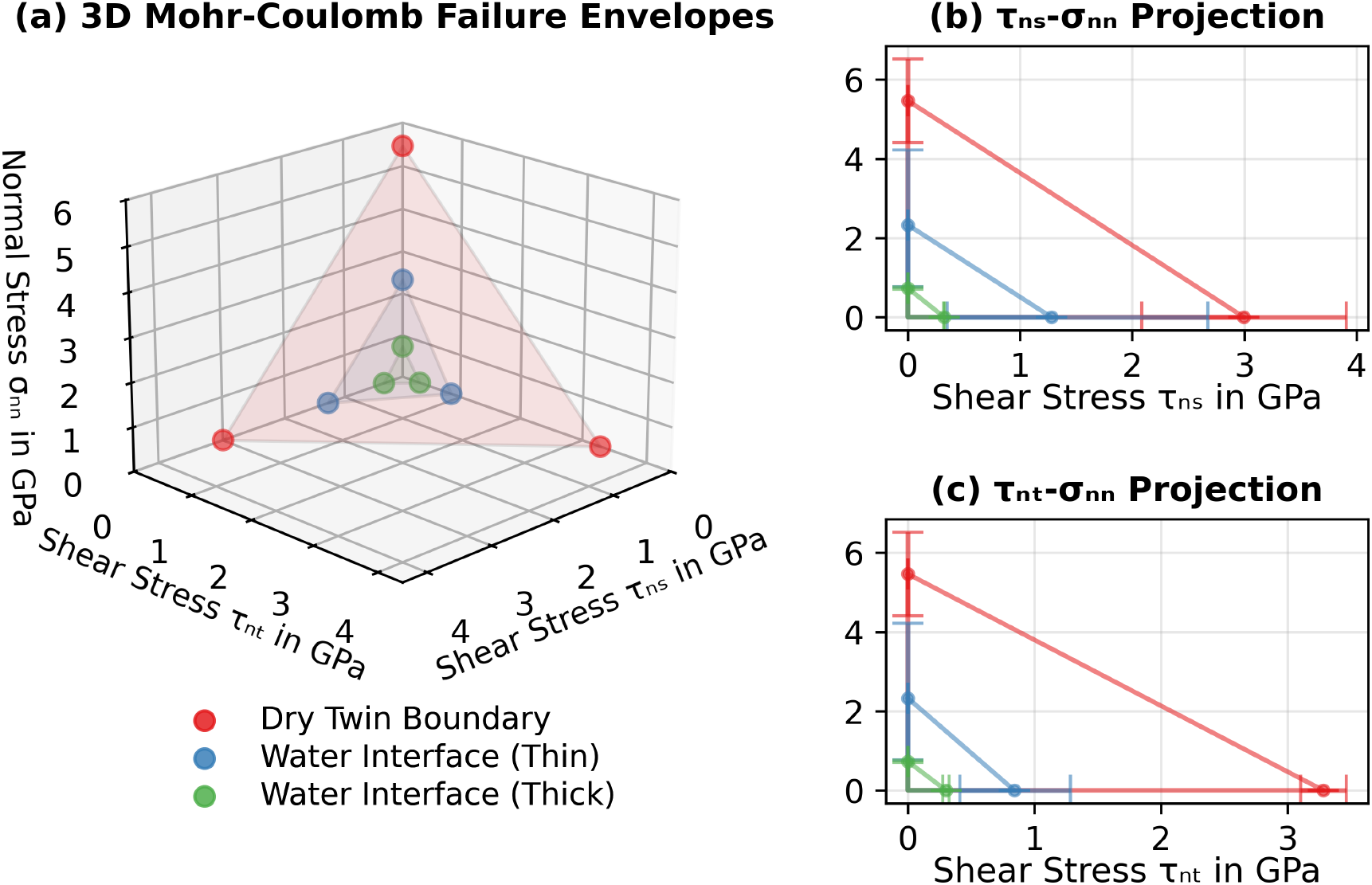
Three-dimensional Mohr–Coulomb failure surface for dry twins and water interfaces. (a)Complete failure surface. (b), (c) Projected stress relationships for quantitative comparison. Bars show strength ranges measured across multiple configurations.

Analysis of failure surfaces (Figure 10) shows distinct performance hierarchies spanning nearly an order of magnitude in strength. Protein-mediated interfaces cluster tightly within a narrow tensile strength range (0.57-0.72 GPa). Single-layer 2JYP interfaces achieve *σ*_*nn*_ = 0.626 *±* 0.050 GPa, with nearly isotropic shear efficiency (*τ*_*ns*_*/σ*_*nn*_ = 0.611, *τ*_*nt*_*/σ*_*nn*_ = 0.574) and low anisotropy (0.06). In contrast, 1L3Q interfaces exhibit lower strength (*σ*_*nn*_ = 0.565 GPa) and higher anisotropy (0.192), potentially reflecting their directional hydrogen-bonded networks and lower water-mediated adhesion observed in Section 3.2.

Two-layer protein interfaces maintain comparable tensile strengths but show intermediate anisotropy (0.088-0.140). This behaviour correlates with the reduced water utilization efficiency and enhanced total binding capacity observed in Section 3.2.

Non-protein interfaces span a much broader strength range (0.73-5.47 GPa, Figure 11). Dry twin boundaries reach *σ*_*nn*_ = 5.47 *±* 1.49 GPa with moderate anisotropy (0.088) and balanced shear ratios (*τ*_*ns*_*/σ*_*nn*_ = 0.55, *τ*_*nt*_*/σ*_*nn*_ = 0.60), consistent with symmetric crystal-crystal bonding. Water interfaces show pronounced thickness effects. Thin water layers (*σ*_*nn*_ ≈ 2.33 GPa) display strong anisotropy (0.343), potentially due to incomplete electrostatic screening. Thick water layers (*σ*_*nn*_ ≈ 0.73 GPa GPa) exhibit reduced anisotropy (0.071) and shear-to-tensile ratios similar to protein interfaces (0.4-0.5).

These results show that interface strength and anisotropy correlate with the balance between electrostatic coupling and hydrogen-bond-mediated adhesion. Direct crystalline contact produces the highest and most isotropic strength, while hydrated interfaces produce lower strength with greater mechanical compliance and energy dissipation capacity.

## 4. Discussion

This study elucidates how the exceptional strength of single-crystal aragonite transitions to the reduced yet resilient mechanical performance of coral skeletons. Through Molecular Dynamics simulations of single crystals and limiting interface compositions — dry twin boundaries, protein-mediated interfaces, and water interfaces — we quantify the mechanical response under tension and shear, establishing clear relationships between interface composition, binding strength, and mechanical performance. We determine that interface chemistry and architecture control this transition. Specifically, protein-aragonite interfaces achieve remarkable damage tolerance because water-mediated hydrogen-bonding networks enable progressive rather than catastrophic failure. This mechanism explores how binding mechanisms evolve during mechanical loading, with sequential bond rupture and reformation identified as the key factor controlling interface failure and energy dissipation. The resulting computational models capture anisotropic interface behaviour across nearly an order of magnitude in strength and provides a quantitative foundation for hierarchical mechanical modelling of biomineral composites.

### 4.1. From Single Crystals to Interfaces

Aragonite’s anisotropic mechanical response originates from distinct atomic-scale deformation mechanisms along its crystallographic directions. Tensile loading along the [100] axis induces deformation-band formation followed by solid-state phase transformation that accommodates strain while maintaining structural integrity. In contrast, [010] loading triggers brittle fracture with minimal post-failure deformation, whereas [001] loading develops organized deformation bands that sustain stress beyond yield. Shear deformation proceeds through coordinated atomic reorganization and band formation rather than bond rupture, producing intrinsic ductility across orientations.

These deformation pathways explain why single-crystal aragonite maintains strengths of 4-6 GPa, consistent with experimental measurements [11, 14–19, 73], while bulk coral skeletons fail already around 0.5 GPa. Introducing nanopores provides additional mechanistic insight: vacancy clusters impede crack propagation and deformation-band development under tension, requiring higher stresses for failure initiation, yet localize shear deformation, thereby reducing shear strength. This contrast illustrates how the same microstructural feature affects strength differently depending on the dominant deformation mode. Neither nanopores nor mesoscale porosity [11, 27, 28] can explain the pronounced softening of coral skeletons, indicating that interfacial processes, rather than crystal defects, govern macroscopic behaviour.

### 4.2. Water-Mediated and Protein-Mineral Adhesion Mechanisms

Interface characterization reveals water-mediated hydrogen bonding as the dominant adhesion mechanism across all simulated configurations. Proteins adhere to aragonite surfaces via structured hydration shells, where bridging water molecules coordinate protein functional groups with surface calcium ions.

The efficiency of these bridging networks depends strongly on interface geometry. Thin interfaces promote dual coordination of water molecules with both protein and mineral, forming dense networks, whereas thicker interfaces reduce this probability, producing more diffuse binding. Consequently, single-layer systems achieve 8.9-17.2% water utilization, whereas two-layer systems reach only 2-3%. This inverse relationship between interface thickness and bridging efficiency reflects geometric constraints of water-mediated adhesion. Bridging water networks preferentially involve protein carboxyl and hydroxyl groups due to their electrostatic affinity for calcium ions, consistent with observations on biomineral interfaces [21–26, 32, 33, 74, 75]. Network flexibility also enables proteins to adapt their conformation while maintaining contact [33, 44, 76], rationalizing the gradual rather than abrupt loss of adhesion under strain.

Surface calcium density further modulates binding modes. The (100) surface provides the strongest direct protein-mineral adhesion (−2259 kcal/mol), whereas the (010) surface, though weaker in direct binding, achieves highest water utilization (17.2%). This apparent discrepancy reveals complementary adhesion strategies: the (100) surface maximizes electrostatic bonding between surface oxygen atoms and positively charged protein residues, while the (010) surface compensates through extensive water-mediated networks with surface calcium ions. The (001) surface exhibits intermediate binding behaviour, balancing both mechanisms.

This dual hierarchy, i.e., water utilization versus binding energy, highlights the interplay between interface geometry and chemistry. Water utilization follows (010) > (100) > (001), whereas direct binding energies follow (100) > (001) > (010). Together they describe how multiple, complementary binding regimes could maintain adhesion under varying chemical and geometric conditions in coral biomineralization. The coexistence of both strategies in our models suggests that coral interfaces may remain functional despite local chemical variability or hydration changes.

### 4.3. Mechanical Performance of Interfaces

Mechanical testing of interfaces under tensile and shear loading demonstrates that interface compsition and crystallographic orientation jointly control mechanical response. Tensile deformation proceeds through successive rupture and reformation of hydrogen-bonded networks, producing oscillatory stress-strain curves where each peak corresponds to partial network reorganization. This progressive bond breaking dissipates energy and prevents catastrophic fracture, explaining the gradual failure behaviour of protein-mediated interfaces. The observed decrease in peak magnitude with continued deformation reflects sequential loss of effective load-bearing contacts until full separation occurs.

In contrast, shear deformation accommodates strain mainly through geometric rearrangement and interface sliding, maintaining similar stress magnitudes across cycles. The distinction between tensile and shear behaviour reflects different deformation pathways available to hydrated protein networks: tensile loading incrementally compromises adhesive bonds, whereas shear enables sustained reconfiguration without total loss of cohesion. The (010) and (001) interfaces show stress increases at high strains as crystal deformation begins contributing to load transfer, while (100) interfaces soften as protein networks detach, transitioning to grain sliding where water-mediated energy dissipation dominates.

Dry twin boundaries and water interfaces establish the mechanical extremes in our simulations. Twins, dominated by ionic and covalent bonding, display pronounced peaks up to 6.5 GPa followed by either brittle failure or relatively smooth plateaus, whereas thick water interfaces deform smoothly at ≈0.8 GPa through hydrogen-bond rearrangement and viscous-like flow. Thin water layers (< 5*Å*) retain partial Coulombic coupling between mineral surfaces, yielding strengths above 3 GPa but reduced ductility. Protein interfaces occupy an intermediate regime balancing strength and ductility, providing efficient energy dissipation among all tested configurations.

A particularly notable observation is the occurrence of shear-coupled grain boundary migration in (310) twins subjected to *xy* shear (in-plane containing the boundary). This phenomenon, well documented in metallic systems [77–79], manifests here as smooth stress-strain response (solid green curve in Figure 7(d)) indicative of boundary motion accommodating shear deformation while maintaining structural coherence. Its appearance in aragonite twins reveals an additional deformation pathway contributing to ductility at mineral-mineral interfaces. This coupling may represent an atomistic analogue of interfacial sliding observed at higher hierarchical levels in coral skeletons [80–82].

These results suggest that mechanical competence of coral skeletons could arise from hierarchical integration of distinct interfacial mechanisms: crystalline bonding in twins provides structural integrity, water-mediated networks supply compliance, and protein layers enable controlled, energy-dissipative failure under complex loading conditions.

### 4.4. Quantitative Failure Hierarchies and Load Transfer Mechanisms

The three-dimensional failure analysis quantitatively links interface chemistry to mechanical competence. The hierarchy of strength among interfaces follows a consistent pattern: dry twin boundaries > thin water interfaces > thick water interfaces ≈ protein interfaces. This order spans nearly one order of magnitude in strength, from up to 6.5 GPa for twins to 0.6-0.8 GPa for protein-mediated interfaces. These differences reflect fundamentally different load-transfer mechanisms: direct crystal-crystal bonding in twins, residual Coulombic coupling in narrow water gaps, and hydrogen-bonded, water-mediated adhesion in protein-containing interfaces.

Dry twins achieve highest strength through direct ionic and covalent continuity between adjacent crystal lattices. Their failure produces minimal energy dissipation but ensures structural rigidity under tensile loading. Thin water interfaces maintain partial electrostatic coupling between aragonite surfaces, providing moderate strength with limited ductility. As interface thickness increases, Coulombic screening progressively decouples opposing surfaces, lowering strength while promoting compliance. This geometric dependence illustrates a potential design principle: even interfaces of identical chemical composition can be tuned mechanically by altering thickness and thus electrostatic coupling efficiency. Such behaviour could enable organisms to modulate skeletal mechanical properties through nanoscale control of organic layer thickness and hydration [83, 84].

Protein interfaces represent the lower bound of interface strength but exhibit high damage tolerance. Their progressive failure is characterized by multistage bond rearrangement and partial reformation rather than sudden separation. The ratio of shear to tensile strength (*τ*_*ns*_*/σ*_*nn*_ or *τ*_*nt*_*/σ*_*nn*_) exceeds 0.5 for most protein-mediated systems, indicating efficient load transfer under multiaxial stress states. This observation suggests that coral skeletons could resist complex environmental stresses where shear and normal components act simultaneously [29, 30, 85]. In contrast, systems dominated by weaker or less directional hydrogen bonding (such as 1L3Q proteins or thick hydrated layers) display lower strength ratios and higher anisotropy, suggesting potential specialization for predominantly tensile accommodation.

Two-layer protein configurations demonstrate that interface thickness influences not only absolute strength but also mechanical response consistency. While their peak tensile strengths remain similar to single-layer interfaces, anisotropy indices decrease (0.09-0.14), reflecting greater directional uniformity. This behaviour arises from reduced efficiency but enhanced redundancy of their hydrogen-bond networks, which allow localized failure without complete debonding, presenting a potentially important feature for maintaining integrity under repeated environmental loads.

### 4.5. Failure Mechanisms and Energy Dissipation Pathways

Interface types differ not only in peak strength but in how they dissipate energy and progress to failure. Protein-mediated interfaces undergo gradual, multi-stage failure. During early deformation (0-5% strain), protein backbones stretch and hydrogen bonds reorganize, enabling strain accommodation without loss of contact. Between 5% and 25% strain, selective amino acid detachment occurs as specific protein-mineral bonds rupture sequentially, corresponding to multiple stress peaks on stress-strain curves. Beyond ~25% strain, residual protein-water networks lose coherence, leading to full separation. This progressive mechanism explains the characteristic oscillatory mechanical response and non-zero binding energy retention observed in Figure 8. It further indicates that energy dissipation is distributed over the deformation history rather than concentrated at a single event.

Water interfaces dissipate energy through different pathways. Thin water gaps preserve significant Coulombic coupling and support high stresses until screening fails, at which point grain separation occurs. Thick hydrated layers instead accommodate deformation via viscous-like sliding and dynamic hydrogen-bond rearrangement, producing lower peak strength but large strain capacity. Dry twin boundaries, while providing highest load-bearing capacity, generally dissipate less energy per unit deformation than hydrated protein interfaces; however, twin plasticity (post-yield flow and shear-coupled migration) provides measurable accommodation that should not be characterized as purely brittle.

The smooth evolution of binding energy compared with oscillatory stress response highlights that failure at hydrated protein interfaces occurs through bond reorganization rather than abrupt rupture. Even at large strains (>30%), partial adhesion persists, indicating that hydrogen-bonded networks retain a fraction of their load-bearing function. Per-residue binding analyses reveal that some amino acid-mineral contacts remain active after global interface separation, allowing partial adhesion to persist even at large strains.

These results suggest that toughness in coral skeletons could arise from hierarchical complementarity [86, 87]: crystals provide load capacity, twins and boundary migration provide crystalline-level accommodation, thin water layers carry higher loads, and protein-water networks supply dissipative mechanisms that prevent catastrophic failure.

### 4.6. Biological and Environmental Relevance

The mechanical tunability observed here suggests potential biological control mechanisms. By adjusting organic-layer thickness, protein composition, or local hydration, corals could potentially modulate the balance between stiffness and toughness within their skeletons [1–3, 5, 11]. Thin hydrated layers and ordered protein arrangements could provide high local stiffness, while thicker hydrated regions or multilayer proteins could increase compliance and energy dissipation. The observed trade-off between total binding strength and per-residue efficiency in different proteins (2JYP vs. 1L3Q) suggests that functional specialization in biomineralization and adaptive control of interface properties during growth may be possible.

The identified hierarchy of deformation and failure mechanisms may represent adaptation of coral skeletal interfaces to their mechanical environment. Dry mineral-mineral contacts such as twin boundaries provide stiffness and load-bearing capacity, while hydrated organic interfaces enable energy absorption and redistribution under cyclic or unpredictable loading. This hierarchical organization could yield a biomineral system capable of resisting turbulent flow and mechanical impact in marine environments.

The thickness-dependent behaviour of water layers provides a potentially important control parameter: by regulating hydration degree or organic-layer thickness, corals could locally tailor stiffness and toughness. Similar adaptive modulation has been proposed for molluscan nacre and echinoderm skeletons, where alternating brittle and ductile layers produce macroscopic toughness exceeding that of their constituent crystals [86, 87]. Our findings are consistent with this concept in cold-water corals, linking molecular bonding mechanisms to potential mesoscale mechanical adaptability.

The persistence of interfacial adhesion under large strains in our simulations suggests potential resilience to moderate mechanical perturbations in vivo. However, the strong dependence of protein-mineral adhesion on hydrogen bonding and interfacial water structuring also implies potential vulnerability to changes in ocean chemistry that alter hydration-layer organization or ionic composition [11]. Ocean acidification and reduced carbonate availability could disrupt interfacial water networks and weaken adhesion, ultimately compromising skeletal integrity [11]. These sensitivities underscore the importance of coupling atomistic-scale mechanistic models with environmental chemistry to predict coral resilience under future ocean conditions.

### 4.7. Integration with Larger-Scale Models

The quantified interface properties, including failure surfaces, strength ratios, and binding-energy evolution, provide a foundation for physically informed constitutive modelling of coral skeletons. In cohesive-zone and continuum-scale formulations [88–91], protein-mediated interfaces can be represented by multi-stage softening laws or internal variables that capture bond-count and bond-strength evolution. This approach reflects the sequence of atomistic failure events observed here: initial backbone stretching, progressive residue detachment, and eventual network collapse.

The binding-energy versus strain relationships (Figure 8) define both the shape and energetic scale of these softening laws, enabling parametrization of progressive interface degradation. Integrating these atomistically derived parameters into finite-element frameworks could bridge nanoscale mechanisms with mesoscale mechanical response, enabling predictive multiscale simulations of coral skeleton behaviour under complex loading conditions [11].

### 4.8. Limitations and Outlook

Although we capture key bonding and deformation mechanisms, several limitations remain. First, simulations are constrained to nanometre length scales and extreme strain rates (~10^10^ s^−1^) [67, 68], which differ from natural loading conditions. Second, idealised planar interfaces and simplified protein models neglect complex three-dimensional geometries and chemical heterogeneity inherent to biological skeletons. Consequently, the strength values reported here represent specific limiting cases rather than direct experimental equivalents and may overestimate or underestimate real interface behaviour depending on actual microstructural complexity.

In addition, binding energies were evaluated using the total potential energy between defined atomic groups, capturing the interaction contributions from Coulombic, van der Waals, and hydrogen-bond terms as described by the force field. Entropic effects, especially those arising from constrained interfacial water molecules, were not explicitly included. While this approximation yields consistent relative trends among interface types, a complete thermodynamic description would require inclusion of configurational entropy, for example through free-energy perturbation or thermodynamic integration methods.

Nevertheless, systematic exploration of limiting interface compositions identifies mechanistic principles (orientation-dependent deformation, shear-coupled boundary migration, thickness-tuned water screening, progressive protein failure) that may be transferable across scales. Future research should integrate these atomistic insights with experimentally characterized microstructures [11, 20, 92], employing techniques such as EBSD and TEM to define realistic interface geometries. Coupling such structural data with cohesive-zone formulations or multiscale coarse-grained models [93, 94] could enable predictive simulations of coral skeletal mechanics under realistic environmental conditions.

Finally, the observed interplay between direct protein-mineral bonding and hydration-mediated adhesion provides potential design principles for biomimetic materials. Synthetic composites that reproduce these hierarchical interface architectures could achieve tunable combinations of stiffness, toughness, and strength – properties important for the durability of coral skeletons.

## 5. Conclusions

This study explores the transition from single-crystal aragonite strength to significantly less strong yet resilient polycrystalline material of cold-water coral skeletons. Molecular Dynamics simulations reveal that interfacial chemistry and architecture, rather than crystal defects, control this transition.

Interface composition governs mechanical behaviour across nearly an order of magnitude in strength: dry twins provide stiffness through direct lattice continuity, thin water layers offer partial electrostatic coupling and moderate ductility, and protein-mediated interfaces achieve damage tolerance through progressive hydrogen-bond reorganization. These mechanisms together suggest how coral skeletons could combine rigidity with energy dissipation under complex loading conditions.

The refined aragonite potential reproduces experimental elastic constants within 5%, enabling quantitative prediction of both tensile and shear responses. Systematic exploration of protein architectures, interface thicknesses, and crystallographic orientations provides descriptors, such as strength ratios and binding-energy evolution, that can potentially be implemented in multiscale constitutive modelling.

While limited by nanoscale dimensions and idealized geometries, we identify deformation principles, including shear-coupled boundary migration, water-mediated screening, and multi-stage protein failure, that may prove transferable across scales. Integration of these atomistic insights with experimentally resolved interface geometries via EBSD or TEM could enable predictive mesoscale simulations of coral mechanics.

This work provides, to the best of our knowledge, the first quantitative investigation of interface mechanics in cold-water corals. By identifying molecular-scale mechanisms of coral interface resilience, this work advances understanding of biomineralization strategies and provides potential design principles for biomimetic composites that unite stiffness, toughness, and adaptability.

## CRediT authorship contribution statement

**N. Kvashin:** Writing - original draft, Methodology, Software, Investigation, Formal analysis, Data curation. **A. Ozel:** Writing - review & editing, Conceptualization, Methodology, Software. **U. Wolfram:** Writing - review & editing, Conceptualization, Methodology, Formal analysis, Supervision, Funding acquisition.

## Acknowledgements

The project was supported by Leverhulme Trust (Research Project Grant RPG-2020-215) and the Deutsche Forschungsgemeinschaft (German Research Foundation, DFG WO 1543/2-1)

## Appendix A Aragonite Potential Formulations

The Buckingham potential formulation includes short-range repulsion and long-range attraction terms:

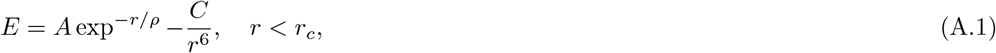

where *A* represents the strength of the repulsive interaction, *ρ* determines the decay rate of the repulsion term, *C* governs the strength of the attractive van der Waals interactions, *r* is the interatomic distance, and *r*_*c*_ is the cut-off radius.

Through our optimization process using GULP, we derive refined coefficients *A, ρ*, and *C* for pair interactions, as well as energy coefficients *K* for intramolecular angle interactions represented by Equations (A.2) and (A.3):

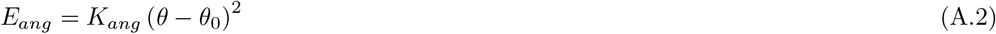

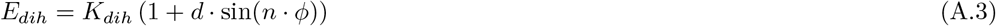

where *θ* and *ϕ* represent bond angles and dihedral angles, respectively, with *θ*_0_ being the equilibrium bond angle. The parameters *K, d*, and *n* control the energetic penalty for deviations from ideal geometry.

## Appendix B Interface Potential

`CHARMM` force fields use Lennard-Jones potentials for description of non-bonded interactions (B.1) and modified terms for bonded interactions (B.2):

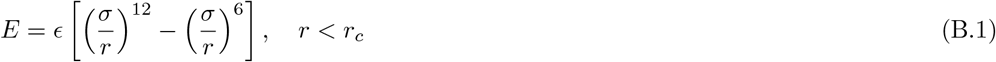

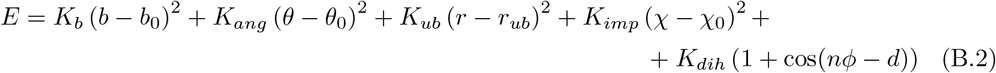

where *r* is the distance between the interacting particles, *ϵ* - the depth of the energy well at the distance 2^1*/*6^*σ, σ* is the distance where the pair interaction energy is zero, *r*_*c*_ is the interaction cut-off; *K*_*b*_, *K*_*ang*_, *K*_*ub*_, *K*_*imp*_, *K*_*dih*_ are the energy constants for bond, angle, Urey-Bradley term for end atoms in an angle triplet, improper, and dihedral interactions within a molecule. The Lennard-Jones part of the potential implemented in `LAMMPS` also includes interactions between outer atoms in consecutively bonded quadruplets.

Next step is the parametrization of interfacial interactions between protein and aragonite constituents which represents a significant challenge in this work. Proteins predominantly consist of hydrogen, oxygen, carbon, and nitrogen atoms, whereas aragonite contains calcium, carbon, and oxygen in different chemical environments. We model interactions between aragonite and protein atoms using a combination of Buckingham potentials (Equation (A.1)) for like atomic species from different constituents and Lennard-Jones potentials from the `CHARMM` force field for dissimilar species (Equation (B.1)).

A crucial challenge lies in parametrizing interactions between calcium from aragonite and the hydrogen and nitrogen atoms in proteins. For these cross-interactions, we apply `CHARMM` force fields using standard Lorentz-Berthelot mixing rules:

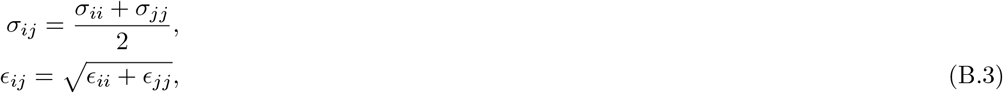

where *σ*_*ii*_ and *ϵ*_*ii*_ are the Lennard-Jones parameters for atomic species *i*, and *σ*_*ij*_ and *ϵ*_*ij*_ are the cross-interaction parameters. The Lennard-Jones parameters for aragonite atoms are derived through fitting using `GULP` to ensure compatibility with the modified Pavese force field.

## Appendix C Mechanical Response of Single Crystal Aragonite

This section provides mechanical property validation supporting the deformation mechanism analysis presented in Section 3.1. The stress-strain behaviour is illustrated in rearrangements that maintain partial load-bearing capacity, producing the stress plateau observed in Figure C.1, showing both tensile and shear responses for loading along the three principal crystallographic directions. Tensile strength values were obtained for loading along the [100], [010], and [001] directions corresponding to *a, b*, and *c* crystal axes, respectively.

**Figure C.1:**
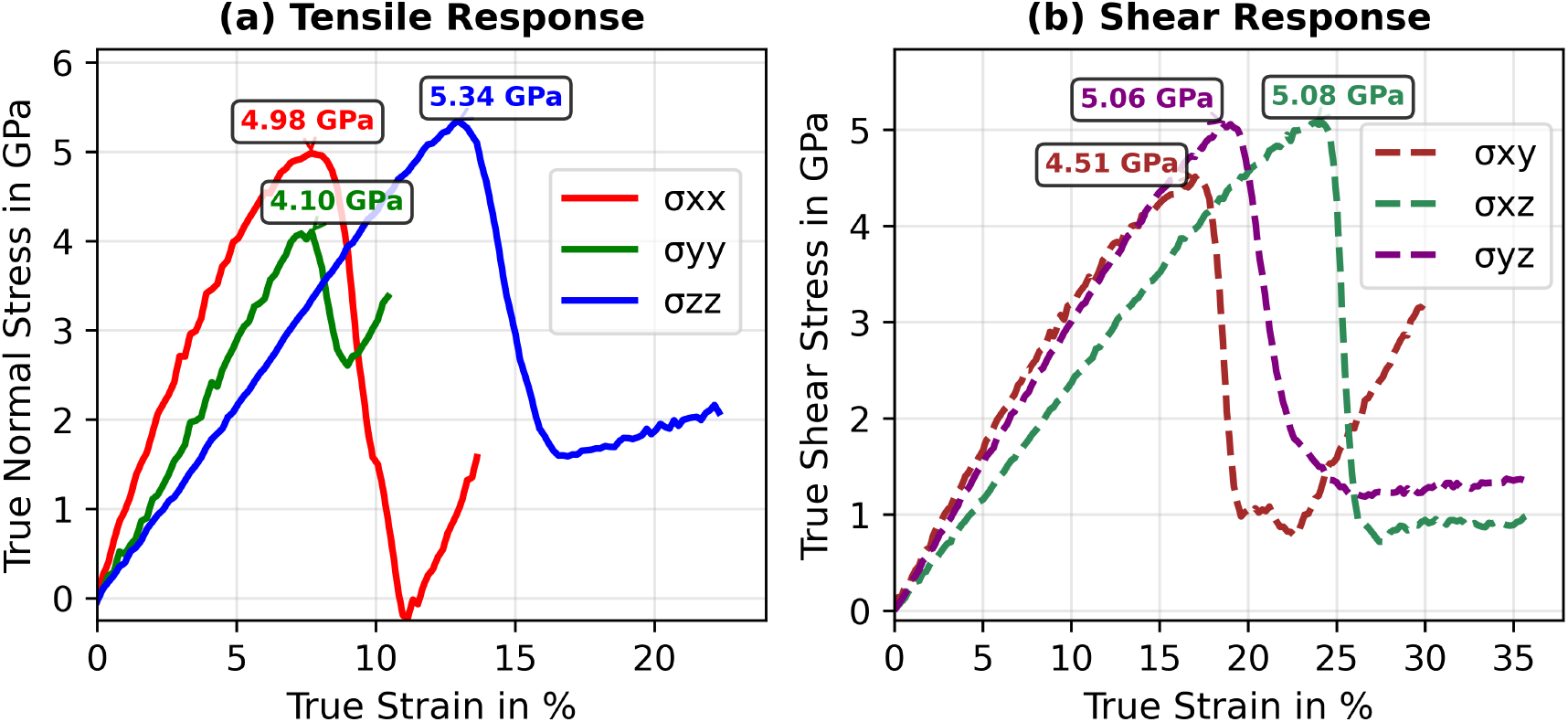
(a) Tensile test of aragonite single crystal showing stress components: *σ*_*xx*_ (red), *σ*_*yy*_ (green), *σ*_*zz*_ (blue) corresponding to [100], [010], and [001] – crystal *a, b*, and *c*axes, respectively. (b) Shear response showing stress components: *σ*_*xy*_ (dark red dashed), *σ*_*xz*_ (dark green dashed), *σ*_*yz*_ (purple dashed)

The [001] direction (along crystal *c*−axis) exhibits the highest strength (5.34 GPa) at 12.9% of applied strain, followed by [100] (4.98 GPa) at 7.7% of applied strain and [010] (4.10 GPa) at 7.7% of applied strain. While directional differences exist, the strength variations (4.10 - 5.34 GPa range) are moderate compared to the scale of property changes observed at interfaces, reflecting the fundamentally crystalline nature of the material with similar bonding environments across orientations.

Shear deformation shows remarkably similar behaviour across the three loading modes (rearrangements that maintain partial load-bearing capacity, producing the stress plateau observed in Figure C.1(b)). All shear responses show peak stresses in the 4.5-5.1 GPa range, with *σ*_*yz*_ and *σ*_*xz*_ exhibiting similar peak values (5.06 and 5.08 GPa respectively) while *σ*_*xy*_ shows slightly lower strength (4.51 GPa). Notably, all shear modes demonstrate exceptional ductility, sustaining deformation to engineering strains exceeding 15% compared to tensile failure. This extended deformation capacity indicates superior load-bearing performance under shear loading conditions. The relatively uniform shear strength across crystallographic orientations indicates that the crystal structure provides consistent resistance to shear deformation regardless of loading direction.

The initial elastic behaviour shows directionally dependent stiffness consistent with the orthorhombic crystal structure (Table 1), with our refined potential accurately reproducing the experimental elastic constants. Beyond the first peak, the simulations reveal complex post-failure behaviour including structural rearrangements and phase transformations as detailed in Section 3.1. However, these high-strain phenomena occur at deformation levels that would be suppressed in real materials by the presence of defects, interfaces, and multi-grain structures that govern failure at much lower strains.

## Appendix D Mechanical Response of (100) Interface Configurations

This appendix provides complete mechanical testing results for all protein configurations in contact with the (100) aragonite surface, which demonstrates the strongest binding energies (Section 3.2). The main text focuses on representative cases spanning the range of interface behaviour across all crystallographic orientations, while this appendix documents the systematic behaviour across all protein architectures on the most energetically favourable surface.

The (100) surface provides the optimal combination of direct protein-mineral interactions and electrostatic compatibility, making it the most relevant for understanding maximum interface performance. All configurations presented here involve loading in the *x*-direction perpendicular to the interfacial plane (crystal *a*-axis) in tensile mode, and *xy*- and *xz*− shear modes within interfacial plane.

### D.1 Tensile Response on (100) Surfaces

The complete tensile testing results for all interface configurations on (100) surfaces reveal systematic trends in mechanical performance related to protein architecture and interface thickness (Figure D.1(a)). Single-layer protein interfaces consistently achieve the highest peak stresses but fail more rapidly, while multi-layer arrangements sacrifice peak strength for enhanced ductility and sustained load-carrying capacity.

Single-layer 1L3Q interfaces demonstrate strength at the first peak around 0.56 GPa. Having superior per-residue binding efficiency (−11.50 kcal/mol per residue) and shorter protein chains, these interfaces demonstrate stable mechanical performance with faster reorganization of protein-water networks under increasing deformation. The relatively smooth stress-strain response without pronounced peaks indicates efficient load transfer through optimized protein-mineral contacts.

Single-layer 2JYP interfaces achieve higher absolute strengths around 0.68 GPa due to larger protein size and greater total binding capacity, but show more complex oscillatory behaviour reflecting the increased complexity of larger protein networks under deformation. The multiple stress peaks indicate sequential failure of different protein domains within the interface.

Both parallel and perpendicular double-layer 2JYP configurations reach peak stresses of 0.70 GPa, representing the intermediate strength among protein-mediated interfaces. The parallel arrangement optimizes protein density while maintaining efficient water-mediated binding networks. The perpendicular protein alignment creates alternative load transfer pathways, although no significant effect on the individual protein-mineral contacts is observed.

Mixed protein interfaces (2JYP+1L3Q) demonstrate the highest strength with peak stresses around 0.72 GPa with unique deformation characteristics reflecting the combined behaviour of both protein types. The heterogeneous protein architecture creates diverse binding mechanisms within the same interface, contributing to sustained load-carrying capacity through sequential activation of different protein-mineral interaction modes. The sustained stress-carrying capacity at high strains demonstrates the effectiveness of all multilayer protein arrangements for energy dissipation.

**Figure D.1:**
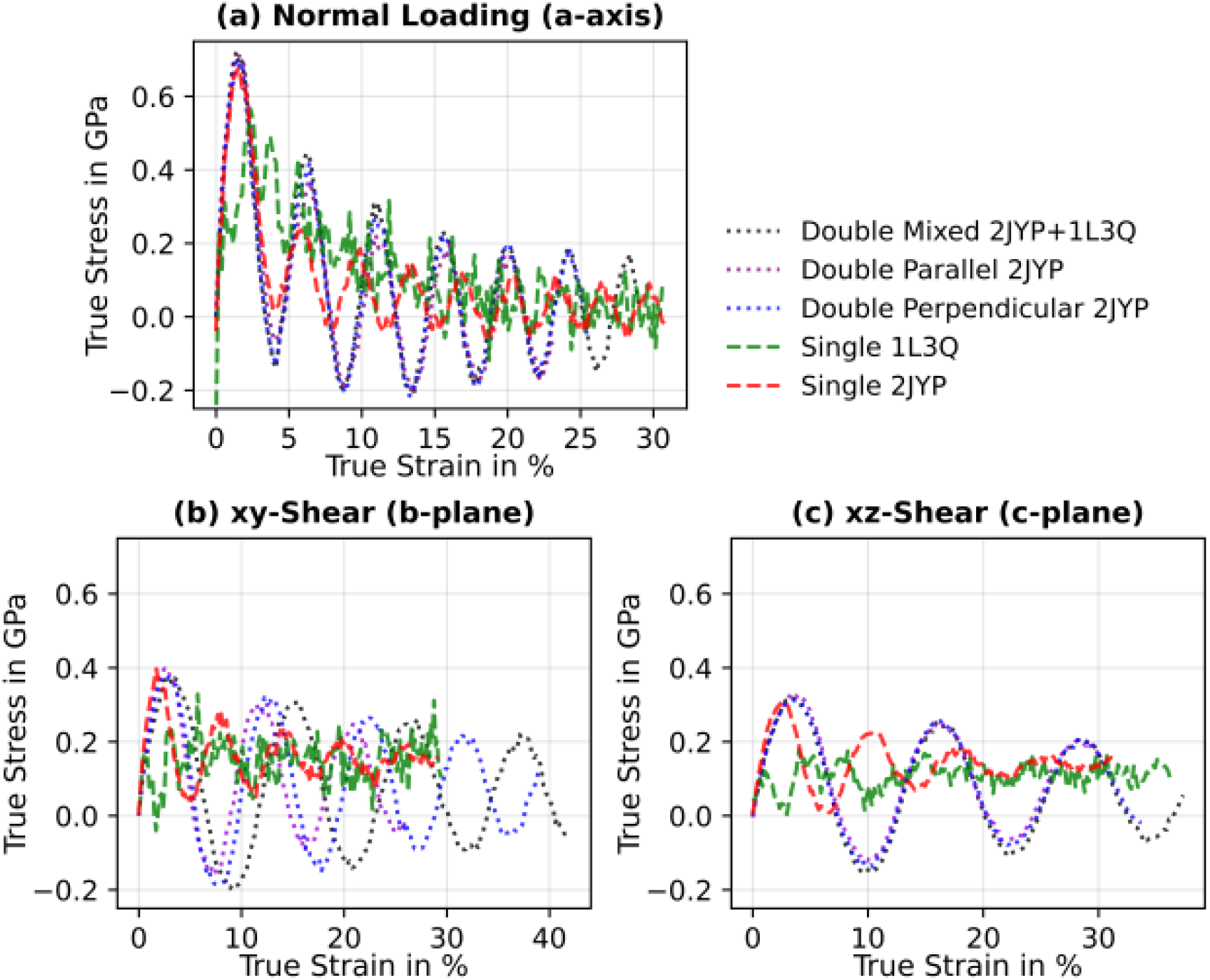
Complete (a) tensile, (b) *xy*− and (c) *xz*− shear testing results for all protein configurations on (100) aragonite surfaces

### D.2 Shear Response on (100) Surfaces

The systematic shear testing of all protein configurations on (100) surfaces reveals consistent accommodation mechanisms across different protein architectures (Figure D.1(b,c)). All protein-mediated interfaces achieve similar peak shear stresses in the 0.3-0.4 GPa range, with variations primarily affecting post-peak behaviour rather than initial strength.

Under *xy*-shear loading, all protein configurations demonstrate similar initial response with peak stresses around 0.4 GPa (Figure D.1(b)). The post-peak behaviour varies systematically with protein architecture: single-layer interfaces show more rapid stress decay, while multi-layer configurations maintain sustained shear resistance through alternative load transfer pathways as individual protein-mineral contacts are progressively compromised.

For *xz*-shear, the protein configurations show consistent behaviour with slightly lower peak stresses around 0.3-0.35 GPa (Figure D.1(c)). The reduced strength in these shear modes may be explained by directional binding differences and protein orientations within the interface. The post yield behaviour is consistent with other shear modes.

The systematic comparison across all protein types confirms that shear accommodation occurs primarily through geometric rearrangement rather than catastrophic bond failure. This fundamental difference from tensile loading explains the sustained performance of protein interfaces under complex stress states and their ability to provide ductile response under marine loading conditions.

